# Machine Learning Identifies Distinct Treg-Mediated Remodeling in HFpEF Hearts Treated with Neonatal Mesenchymal Stem Cells and Their Secretome

**DOI:** 10.64898/2026.01.29.702616

**Authors:** Zhi-Dong Ge, Jonghyeuk Han, Felipe Takaesu, Rachana Mishra, Sudhish Sharma, Ling Chen, Xuebin Fu, Mallory Filipp, Ching Man Wai, Ning-Yi Shao, Anshuman Sinha, Progyaparamita Saha, Akash Puvvala, Luna Ventura, Sonia Thakkar, Agata Bileweska, Artur Stefenwicz, Sameer Ahmad Guru, Muthukumar Gunasekaran, Evan Yang, Buddhadeb Dawn, Peixin Yang, Sanjiv J. Shah, Edward Thorp, Michael E. Davis, Sunjay Kaushal

**Affiliations:** Cardiovascular-Thoracic Surgery and the Heart Center, Stanley Manne Children’s Research Institute, Ann & Robert H. Lurie Children’s Hospital of Chicago, Departments of Pediatrics and Surgery, Feinberg School of Medicine, Northwestern University, 225 E. Chicago Avenue, Chicago, Illinois 60611, USA; Wallace H. Coulter Department of Biomedical Engineering, Georgia Institute of Technology and Emory University School of Medicine, Atlanta, GA, 30332, USA; Biochemistry, Cell and Developmental Biology Graduate Training Program, Graduate Division of Biological and Biomedical Sciences, Laney Graduate School, Emory University, Atlanta, GA, 30332, USA; Department of Pathology, Feinberg School of Medicine, Northwestern University, 300 E. Superior Avenue, Chicago, Illinois 60611, USA; Department of Biochemistry and Molecular Genetics, Feinberg School of Medicine, Northwestern University, 300 E. Superior Avenue, Chicago, Illinois 60611, USA; Faculty of Healthy Sciences, University of Macau, Macau, China; Renaissance School of Medicine, Stony Brook University, 100 Nicolls Road, Stony Brook, NY, 11794, USA; Department of Medicine, Northwestern Memorial Hospital, Feinberg School of Medicine, Northwestern University, 676 N. Saint Clair, Chicago, Illinois 60611, USA; Division of Cardiology, Emory University School of Medicine, Atlanta, GA, 30322, USA; Children’s Heart Research and Outcomes (HeRO) Center, Children’s Healthcare of Atlanta and Emory University, Atlanta, GA, 30322, USA; Department of Obstetrics, Gynecology, and Reproductive Sciences, University of Maryland School of Medicine, Baltimore, Maryland, USA; Department of Internal Medicine, Division of Cardiovascular Medicine, Kirk Kerkorian School of Medicine at the University of Nevada Las Vegas, Las Vegas, NV 89102, USA; Department of Surgery, Kirk Kerkorian School of Medicine at the University of Nevada Las Vegas, Las Vegas, NV 89102, USA

**Keywords:** Heart failure with preserved ejection fraction (HFpEF), single-nucleus RNA sequencing, causal discovery, machine learning, nMSCs, secretome, intercellular communication, cardiac remodeling, T_reg_ cells, PLS regression

## Abstract

**Background:** Heart failure with preserved ejection fraction (HFpEF) remains a major therapeutic challenge due to its complex pathophysiology and pronounced heterogeneity. Regenerative approaches using neonatal mesenchymal stromal cells (nMSCs) and their secretome (SEC) have shown promise in other heart failure contexts.

**Objectives:** However, the effect of these therapies in HFpEF, and the underlying molecular mechanisms and causal pathways remain poorly understood.

**Methods:** HFpEF was established in two distinct murine models, followed by treatment with either nMSCs or SEC. Functional and histological endpoints were assessed. We developed a novel machine learning framework, VIPcell, which integrates data augmentation, Partial Least Squares (PLS) regression, and causal structure inference to identify genes causally linked to cardiac function using single-nucleus RNA sequencing (snRNA-seq) data. VIPcell was applied to heart tissues from treated HFpEF animals to uncover key regulators of cardiac remodeling.

**Results:** Both nMSC and SEC therapies significantly improved diastolic function in two independent rodent HFpEF models. These improvements were associated with reduced inflammation, attenuated myocardial fibrosis, and improved exercise capacity. Intercellular communication analysis revealed widespread, system-level signaling in nMSC-treated hearts, compared to more localized endothelial–cardiomyocyte crosstalk in SEC-treated hearts. Causal inference via VIPcell suggested overlapping upstream regulators in both treatment groups, particularly genes involved in regulatory T cell (T_reg_) biology and immunomodulatory signaling pathways, including FOXO signaling, NLRP3 inflammasome inhibition, and Tie2 activation. In vivo validation confirmed selective expansion of T_regs_ following nMSC and SEC therapy. In vitro, nMSCs induced significantly greater T_reg_ expansion compared to multiple adult stem cell types. Critically, chemical depletion of T_regs_ abrogated the therapeutic effects of both treatments, establishing T_regs_ as central mediators of diastolic function recovery in the HFpEF preclinical model.

**Conclusions:** nMSC and SEC therapies improve diastolic function in HFpEF through distinct remodeling mechanisms converging on T_reg_-mediated immune modulation. VIPcell supported identification of causal regulators, highlighting T_reg_-related signaling as a key driver of myocardial recovery in HFpEF. These findings offer mechanistic insight into cellular therapies for HFpEF and support the development of targeted, T_reg_-focused interventions.

## Introduction

Heart failure with preserved ejection fraction (HFpEF) represents a rapidly escalating clinical and public health epidemic, now accounting for at least half of all heart failure diagnoses worldwide.^1,2^ Despite its growing prevalence and the recent advent of multiple evidence-based treatments, HFpEF remains a therapeutic challenge with high residual risk of HF hospitalization and cardiovascular death. In contrast to heart failure with reduced ejection fraction (HFrEF), HFpEF is characterized by predominant impairment in LV diastolic relaxation and compliance with varying degrees of subtle systolic dysfunction with preservation of global LV ejection fraction, often arising in the context of comorbidities such as obesity, hypertension, diabetes, and systemic inflammation.^3–5^ These multifactorial conditions initiate a cascade of pathophysiologic changes—including myocardial stiffness, microvascular dysfunction, chronic inflammation, immune dysregulation, and interstitial fibrosis—that culminate in reduced ventricular compliance and exercise intolerance.^3,6,7^ The clinical heterogeneity and mechanistic complexity of HFpEF have contributed to the limited success of traditional pharmacologic therapies, underscoring the urgent need for novel, mechanistically informed treatment strategies.^8^

Therapeutic approaches leveraging mesenchymal stromal cells (MSCs) and their paracrine derivatives—collectively referred to as the secretome (SEC)—have emerged as promising alternatives to address the underlying drivers of cardiac dysfunction.^9–11^ These therapies exert their therapeutic effects predominantly through paracrine signaling, modulating inflammation, enhancing angiogenesis, reducing oxidative stress, and orchestrating tissue repair through immune-mediated pathways. Neonatal-derived MSCs (nMSCs), in particular, exhibit enhanced reparative potential relative to adult-derived counterparts, due their robust immunomodulatory capacity, higher proliferative activity, and enriched secretion of pro-regenerative factors.^12–19^ Human nMSCs possess the unique ability to evade the immune response and proliferate within the injured myocardium in many tested rodent and porcine animal models - critical for their robust efficacy as an allogenic preparation needed for future stem cell trials. The nMSC-derived secretome (SEC), composed of cytokines, growth factors, coding and non-coding RNAs, and extra vesicles, recapitulates many of the therapeutic benefits of cell-based therapy while offering advantages in scalability, production cost, and ease of use.^18^ nMSCs and their SEC have shown efficacy across a broad range of preclinical models which are driven by acute and/or chronic inflammation—including rodent and swine models of HFrEF, acute kidney injury, amyotrophic lateral sclerosis (ALS), Alzheimer’s disease, chronic ileitis, cardiopulmonary bypass-induced systemic inflammation, and right ventricular pressure overload.^15,18,20–23^ However, despite these promising findings, the molecular mechanisms through which nMSCs and SEC mediate cardiac repair remain incompletely understood—particularly in HFpEF, where immune dysregulation and endothelial dysfunction play central roles in disease progression and therapeutic resistance.

To address this critical knowledge gap, we developed VIPcell (Variable Importance Prediction for cellular function), a novel machine learning framework designed to infer causal gene-function relationships from single-nucleus RNA sequencing (snRNA-seq) data. VIPcell integrates data augmentation, Partial Least Squares (PLS) regression for variable importance ranking, and causal structure inference using a Discovery at Scale (DAS) algorithm to identify genes that causally influence tissue-level phenotypes. Applying VIPcell to snRNA-seq data from HFpEF mouse models treated with either nMSCs or their SEC, we aimed to delineate shared and distinct cellular remodeling mechanisms contributing to improved diastolic function. This study provides three key advances: (1) a scalable computational pipeline for causal gene discovery in limited number of high-dimensional single-cell data; (2) a systems-level comparison of the therapeutic remodeling effects of nMSCs versus SEC in HFpEF; and (3) mechanistic insight into the role of T_reg_ and immune-mediated repair pathways—including FOXO signaling, Tie2 activation, and NLRP3 inflammasome modulation—in mediating cardiac recovery. Collectively, these findings not only advance our understanding of the remodeling logic underlying nMSC and SEC therapies but also establish a platform for mechanistically guided therapeutic optimization in HFpEF and related conditions.

## RESULTS

### Both nMSC and SEC improve cardiac diastolic function in HFpEF models

The majority of patients with HFpEF have increased visceral adiposity, hypertension, metabolic syndrome, and frequently have diabetes and pulmonary congestion.^43^ To model this cardiometabolic phenotype, C57BL/6 mice were fed a HFD in combination with L-NAME, a constitutive nitric oxide synthase inhibitor, as previously described.^44^ This approach induces metabolic and nitrosative stress, along with hypertension, collectively modeling HFpEF pathophysiology. Mice were fed HFD with L-NAME for five weeks to induce HFpEF and continued to have this diet throughout the remaining study and were randomized to receive normal saline placebo (HFpEF), SEC, or nMSCs. A control group received a normal chow diet throughout the study (Figure 1A, Supplement Fig.1). Cardiac structure and function were assessed via echocardiography. nMSCs and their SEC were isolated, expanded, and characterized as previously described^15,18,20–23^.

**Figure 1.**
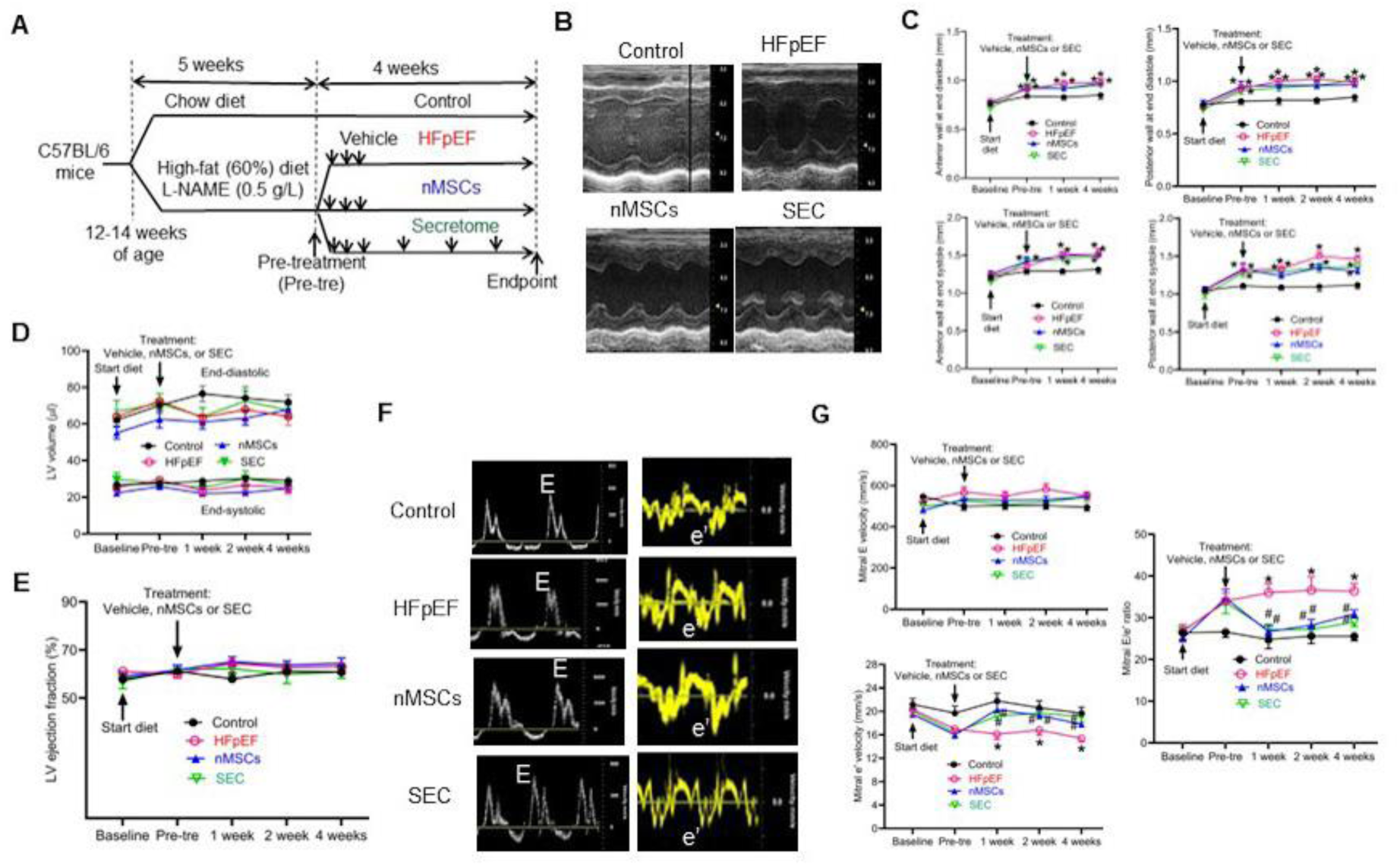
Both nMSCs and SEC improved left ventricular diastolic function but did not affect systolic function in the mice of heart failure with HFpEF. A: Experimental procedures. L-NAME, L-N^G^-nitroarginine methyl ester. B: Representative echocardiographs of M-mode images of the left ventricle at the endpoint of the experiments showing left ventricular hypertrophy in the mice of HFpEF, nMSCs, and SEC groups. C: Changes in the wall thickness of the left ventricle in the mice of control, HFpEF, nMSCs, and SEC groups (n = 12-14 mice/group) throughout the experiments. D: Changes in left ventricular (LV) volumes at end diastole (Top panels) and at end systole (Bottom panels) in the mice of control, HFpEF, nMSCs, and SEC groups (n = 12-14 mice/group) throughout the experiments. E: Changes in LV ejection fraction in the mice of control, HFpEF, nMSCs, and SEC groups (n = 12-14 mice/group) throughout the experiments. F: Represative echocardiographs of pulsed Doppler-mode images of mitral waves (Left) at the endpoint of the experiment showing mitral E waves and tissue Doppler-mode images (Right) showing mitral e’ waves in 4 groups of mice. G: Changes in mitral E and e’ waves in 4 groups of mice throughout the experiments (n = 12-14 mice/group). *P<0.05 versus control groups, and ^#^P < 0.05 versus HFpEF groups.

We first evaluated left ventricular (LV) wall thickness, volume, and ejection fraction (EF) using short-axis B-mode-guided M-mode imaging (Figure 1B). At baseline, there were no significant differences in these parameters across the experimental groups. At both pretreatment and endpoint timepoints, HFpEF mice demonstrated LV hypertrophy without significant alterations in LV volumes or EF compared to controls. Furthermore, no significant differences were observed in anterior/posterior wall thickness at end-diastole and end-systole, LV volumes or EF among the HFpEF, nMSC, and SEC treatment groups (Figures 1C–1E). We used WGA to stain mouse heart sections and quantified cardiomyocyte size. Cardiomyocyte surface area was larger in HFpEF than control groups (Supplement Figure 8). There were no significant differences in cardiomyocyte surface area among nMSCs, SEC, and HFpEF groups. These results indicate that nMSC and SEC treatment did not reverse cardiac hypertrophy or affect LV systolic function in HFpEF mice.

To assess diastolic function, we measured peak mitral E and e′ wave velocities using apical four-chamber view-guided pulsed-wave Doppler and tissue Doppler imaging of the mitral annulus (Figure 1F, Supplement Figure1). At baseline, mitral E and e′ velocities, as well as the E/e′ ratio, were similar across all groups (Figure 1G). During the study, mitral E velocity remained unchanged. However, at the pretreatment timepoint, e′ velocity was reduced, and E/e′ ratio was elevated in the HFpEF, nMSC, and SEC groups, consistent with impaired diastolic function. Importantly, by the endpoint, e′ velocity improved, and the E/e′ ratio decreased in both the nMSC and SEC groups relative to placebo-treated HFpEF mice. These findings suggest that nMSC and SEC therapy both improve diastolic function (particularly early diastolic relaxation) in the HFpEF model, without affecting systolic performance or cardiac remodeling.

To validate these findings in a second HFpEF model and using equal number of male and females, we utilized the leptin receptor-deficient (db/db) mouse, a well-established model of obesity and Type 2 diabetes, alongside nondiabetic heterozygous db/+ littermates as controls. Diastolic dysfunction in db/db mice was confirmed by an elevated E/e′ ratio, which emerged by 12 weeks of age and plateaued by 24 weeks (Supplement Figure 3,4). Db/db mice were then randomized to receive either nMSC, SEC, or placebo, while db/+ mice served as baseline controls at 24 weeks. Following treatment, both nMSC and SEC significantly improved diastolic function, as evidenced by e′ velocity improvement with no change in E velocity and the improvement of E/e′ ratio in both the nMSC and SEC groups relative to placebo-treated HFpEF mice reduced E/e′ ratios (Supplement Figure 5). These improvements occurred without changes in anterior/posterior wall LV thickness (Supplement Figure 6). Together, these findings demonstrate that both nMSC and SEC therapies consistently rescue diastolic dysfunction across two distinct murine models of HFpEF, independent of effects on systolic function or cardiac hypertrophy.

### Both nMSC and SEC improve general physiological function and cardiac fibrosis in HFpEF

Decreased exercise intolerance is a defining clinical feature of heart failure, including HFpEF.^45^ To evaluate this, treadmill testing was conducted at the study endpoint to assess exercise capacity. The HFpEF group in both HFpEF models of HFD/L-NAME or db/db demonstrated significantly reduced running distance, speed, and duration compared to controls (Figure 2A, Supplement Figure 6). Notably, treatment with nMSCs led to improvements in running distance and speed but not running time, while treatment with SEC caused improvements in all three parameters at the 4-week timepoint (Figure 2A, Supplement Figure 7). These findings suggest that nMSC and SEC therapies improve overall physiological performance in different HFpEF models, potentially translating into enhanced quality of life in clinical settings.

**Figure 2.**
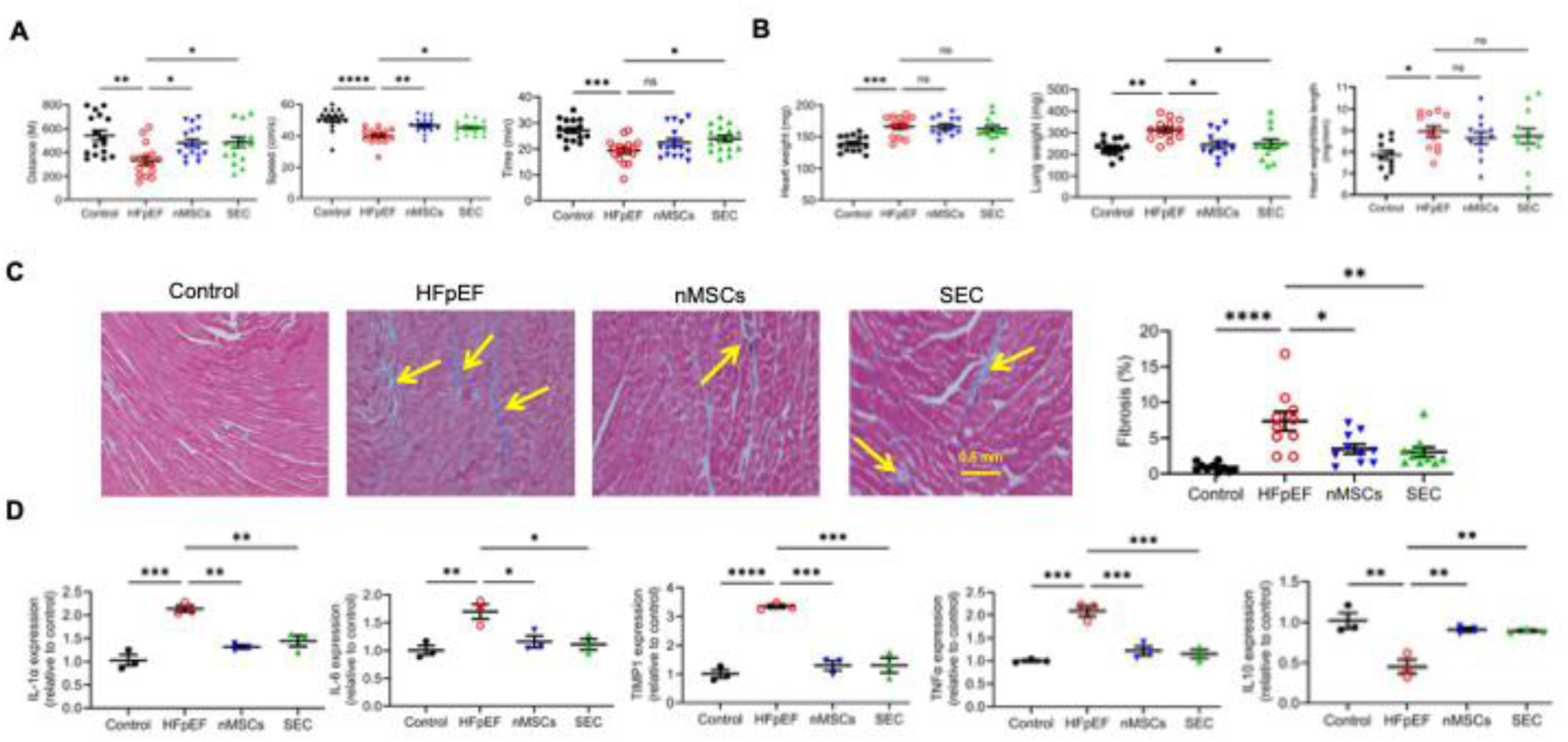
Both nMSCs and SEC improved general physiological function, prevented myocardial fibrosis, and regulated pro-inflammatory cytokines in the mice of HFpEF. A: Distance, speed, and time measured on treadmill exercise stress test in the mice of control, HFpEF, nMSCs, and SEC groups at the endpoint of the experiment (n = 16 mice/group). Mice were run for 5 min at a speed of 5 m/min, then for 2 min at 14 m/min. The speed was increased by 2 m/min every 2 min until mice reached exhaustion. B: Wet heart weight, lung weight, and the ratio of heart weight/tibia length in the mice of 4 groups at the endpoint of the experiment (n = 11-13 mice/group). C: Myocardial fibrosis in the mice of 4 groups at the endpoint of the experiment (n = 10 heart sections in 3 mice/group). Paraffin-fixed mouse heart sections were stained with Masson’s trichrome. The arrowheads point to fibrosis. Scar bar: 0.5 mm. D: Changes in pro-inflammatory cytokines in the mice of 4 groups at the endpoint of the experiment (n = 3 mice/group). IL-1α, interleukin 1α; IL-6, interleukin 6; TIMP1, TIMP metallopeptidase inhibitor 1; TNFα, tumor necrosis factor-alpha; IL-10, interleukin 10. *P<0.05 versus control groups, and ^#^P < 0.05 versus HFpEF groups.

After euthanasia, we recorded body, heart, and lung weights in the HFD/L-NAME model of HFpEF. Body weight was significantly higher in the HFpEF, nMSC, and SEC groups compared to placebo, with no significant differences among the treated and untreated HFpEF groups. In contrast, heart and lung weights were significantly elevated in HFpEF mice versus placebo, reflecting cardiac hypertrophy and pulmonary congestion. The heart weight-to-tibia length ratio, a surrogate for cardiac remodeling, was similarly increased in the HFpEF group. However, no significant differences in heart weight or heart weight-to-tibia length ratio were observed between nMSC-, SEC-treated, and untreated HFpEF groups (Figure 2B).

Cardiac fibrosis is a near-universal feature in symptomatic HFpEF and can contribute to myocardial stiffening and diastolic dysfunction.^46,47^ To assess myocardial fibrosis, Masson’s trichrome staining was performed on harvested heart tissue. As expected, fibrosis was significantly elevated in the HFpEF group compared to controls in the HFpEF db/db model (Figure 2C). Importantly, both nMSC and SEC treatment resulted in a significant reduction in myocardial fibrosis relative to untreated HFpEF mice (Figure 2C). These results indicate that nMSC and SEC therapies attenuate cardiac fibrosis, a key pathologic hallmark of HFpEF, supporting their potential to reverse structural remodeling in this disease model.

Cytokines are critical regulators of the inflammatory response, which is a central contributor to the pathogenesis of HFpEF.^48–51^ To investigate the anti-inflammatory effects of nMSCs and SEC, we assessed the cardiac gene expression of key inflammatory mediators, including interleukin (IL)-1α, IL-6, tumor necrosis factor-α (TNF-α), tissue inhibitor of metalloproteinase 1 (TIMP1), and the anti-inflammatory cytokine IL-10, in HFpEF HFD/L-NAME mouse hearts. As shown in Figure 2D, compared to control animals, HFpEF mice exhibited significantly increased expression of IL-1α, IL-6, TIMP1, and TNF-α, along with decreased expression of IL-10, consistent with a pro-inflammatory cardiac environment. Notably, treatment with nMSCs or SEC resulted in a significant downregulation of IL-1α, IL-6, TIMP1, and TNF-α expression compared to the untreated HFpEF group (Figure 2D). Additionally, IL-10 expression was significantly upregulated in both treatment groups relative to HFpEF controls (Figure 2D). These findings indicate that nMSC and SEC therapies modulate the inflammatory milieu in HFpEF by suppressing pro-inflammatory cytokine expression while enhancing anti-inflammatory signaling, supporting their immunoregulatory and therapeutic potential in chronic inflammatory cardiac conditions.

### Cellular and Interaction Atlas after nMSC and SEC Treatment

To further determine the molecular mechanism central to their recovery, we performed snRNA-seq on a total of 26,410 nuclei isolated from adult C57BL/6 mice hearts across four experimental groups: control, HFpEF, nMSC, and SEC in the HFD/L-NAME model of HFpEF. Unsupervised clustering identified 19 transcriptionally distinct cell clusters, which were visualized in two-dimensional (2D) UMAP space (Figure 3A). To annotate each cluster, we cross-referenced the top differentially expressed genes with curated cell type markers from publicly available databases, including CellMarker 2, PanglaoDB, and CZ CELLxGENE. This analysis identified a diverse cellular landscape within the heart, comprising cardiomyocytes, fibroblasts, endothelial cells, macrophages, mural cells, endocardial cells, erythroid-lineage cells, sinoatrial nodal cells, mesothelial cells, lymphocytes, smooth muscle cells, pericytes, lymphatic endothelial cells, and glial cells. Hierarchical clustering, visualized through a heatmap, revealed distinct gene expression profiles across all identified cell types. Notably, we observed transcriptional heterogeneity within the two most abundant populations: cardiomyocytes and endothelial cells. Cardiomyocytes emerged as the predominant population and were further subdivided into four transcriptionally distinct subpopulations, each expressing canonical cardiomyocyte markers including *Fhod3*, *Rbm20*, *Corin*, and *Mhrt* (Figure 3B). Among these, cardiomyocyte subpopulation III was enriched for *Ankrd1*, *Ankrd23*, and *Nppb*, genes commonly upregulated in heart failure and myocardial stress responses. In contrast, subpopulation IV was characterized by expression of *Nppa* and *Myl4*, markers indicative of atrial cardiomyocytes. Endothelial cells constituted the second largest population, comprising three subclusters defined by shared markers *Pecam1*, *Flt1*, and *Egfl7*. Interestingly, each endothelial subcluster also expressed a unique distinguishing gene: *Flt1* (endothelial cell I), *Tmsb4x* (endothelial cell II), and *Cyyr1* (endothelial cell III). These genes have not been extensively studied in the context of endothelial cell heterogeneity, highlighting novel molecular distinctions and potential functional specialization within the endothelial compartment.

**Figure 3.**
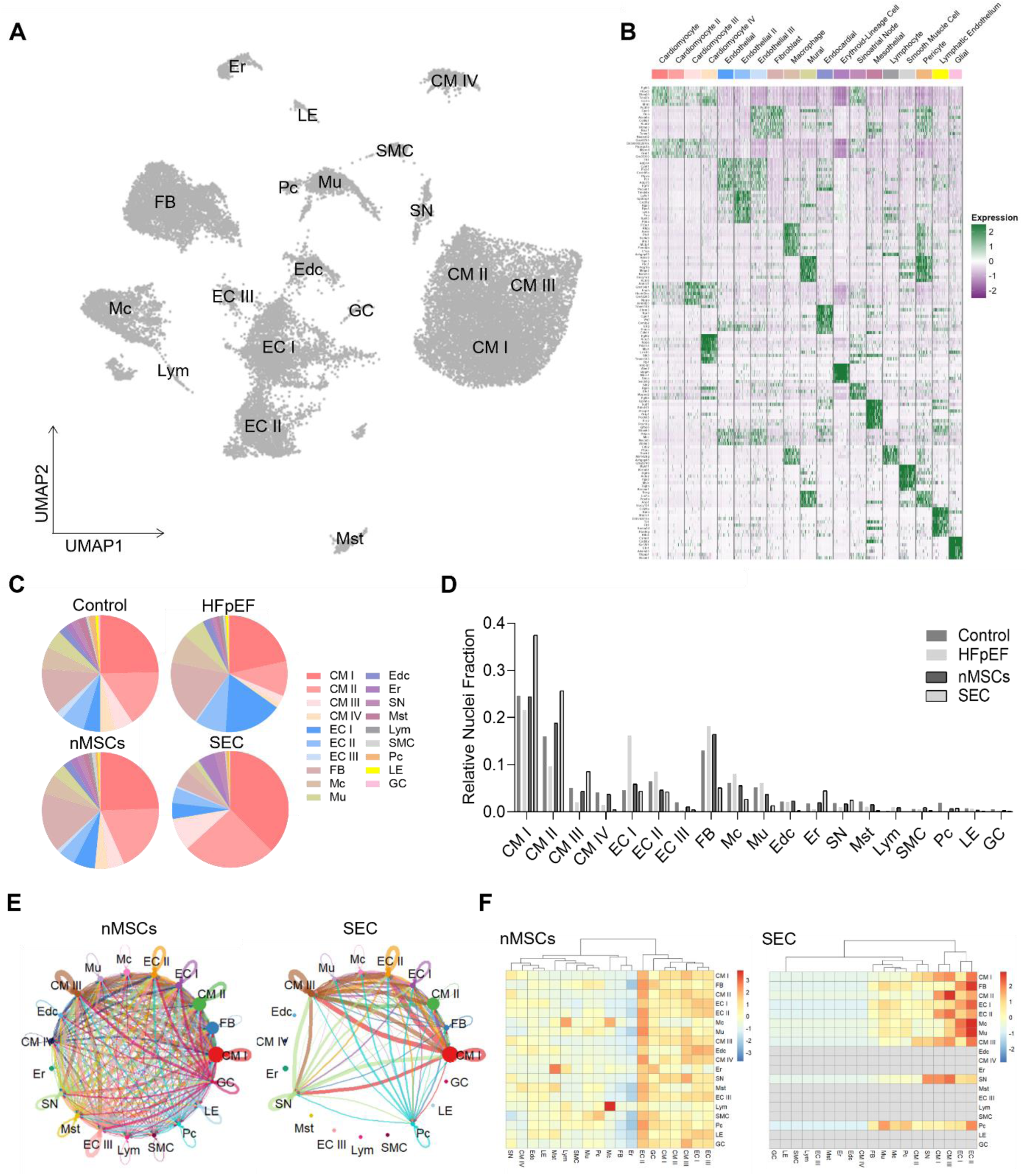
nMSCs and SEC treatment induced distinct cellular and cell-cell signaling atlas in tissues. A: UMAP projection of clusters formed by 26,410 cardiac cells. B: The heatmap of top differentially expressed genes in each cell clusters. C: Pie chart depicting the relative nuclei fraction of each cell clusters for Control, HFpEF, nMSCs- and SEC-treated tissues. The number of nuclei in each cluster is normalized by the total number of nuclei in the tissue. D: Graph showing the relative nuclei fraction in WT/ HFpEF/ nMSCs/ SEC groups. E: Circle plot and f, Heat map showing strength of significant outgoing interactions from source cell cluster to target clusters inferred by CellChatDB. (Left: nMSC, Right: TCM) For circle plot, each circle represents a cell cluster. The size of the circle reflects the cluster fraction. The line width means the strength of interaction and each color means the outgoing interaction from the source cluster. For heatmap, Colors represent the strength of significant outgoing interactions from a source cell cluster (rows) to target clusters (columns). CM: cardiomyocyte, FB: fibroblast, EC: endothelial cell, Mc: macrophage, Mu: mural cell, Edc: endocardial cell, Er: erythroid-lineage cell, SN: sinoatrial node, Mst: mesothelial cell, Lym: lymphocyte, SMC: smooth muscle cell, Pc: pericyte, LE: lymphatic endothelium, GC: glial cell

We next compared cellular heterogeneity among the control, HFpEF, nMSC-treated, and SEC-treated heart tissues by analyzing the nuclei fraction of each cell cluster within each group (Figure 3C–D). In the control group, cardiomyocytes accounted for approximately 50% of the total nuclei, followed by endothelial cells (13.25%), fibroblasts (13%), and macrophages (6.25%). In the HFpEF group, there was a marked increase in endothelial cell subcluster I, along with elevated proportions of fibroblasts and macrophages, consistent with the inflammatory and fibrotic remodeling seen in HFpEF. Interestingly, the nMSC- and SEC-treated groups displayed distinct patterns of cellular regeneration. The nMSC-treated hearts exhibited a cellular composition closely resembling that of the healthy control group, suggesting a restorative effect. In contrast, the SEC-treated hearts showed a substantial increase in the proportion of cardiomyocytes to 72.4%, with enrichment specifically in non-atrial cardiomyocyte subpopulations I, II, and III, indicating a potentially enhanced remodeling effect on myocardial cell populations.

Given that nMSC and SEC treatments reshape the cellular landscape in HFpEF heart tissue, we next examined intercellular communication networks across cell clusters using the CellChat R package. In the nMSC-treated group, signaling interactions were broadly distributed across diverse cell types, indicating a global pattern of intercellular communication. In contrast, the SEC-treated group exhibited a more restricted distribution, with signaling interactions concentrated primarily among endothelial cell and cardiomyocyte subpopulations (Figure 3E). These distinct communication profiles were further illustrated in heatmaps of incoming and outgoing signaling activity (Figure 3F). In the nMSC group, nearly all clusters exhibited both incoming and outgoing signaling, underscoring a multicellular and coordinated network. In the SEC group, however, outgoing signals were predominantly derived from cardiomyocytes, endothelial cells, macrophages, mural cells, sinoatrial nodal cells, and pericytes, whereas incoming signals were primarily directed toward cardiomyocyte subpopulations I and III and endothelial subpopulations I and II. These findings suggest that intercellular communication is particularly enriched between cardiomyocytes and endothelial cells in the SEC-treated hearts, consistent with the increased abundance of these cell types. Collectively, our results indicate that nMSC and SEC therapies elicit distinct remodeling mechanisms: nMSCs promote a broadly distributed signaling network, whereas SEC treatment engages a more targeted and focused communication axis, particularly between cardiomyocytes and endothelial cells.

### Unbiased Identification of Causal Genes for HFpEF Recovery via a VIPcell Framework

To further investigate the distinct remodeling mechanisms underlying nMSC- and SEC-treated HFpEF hearts, we developed a novel machine learning (ML)-based framework to identify key molecular drivers of tissue function from single-nucleus RNA sequencing data. The VIPcell framework is step-by-step process, integrating 1) scRNA-seq augmentation and conversion, 2) Partial Least Squares (PLS) regression to identify high-importance genes associated with function, and 3) causal inference via the DAS to distinguish causality from correlation (Figure 4A).

**Figure 4.**
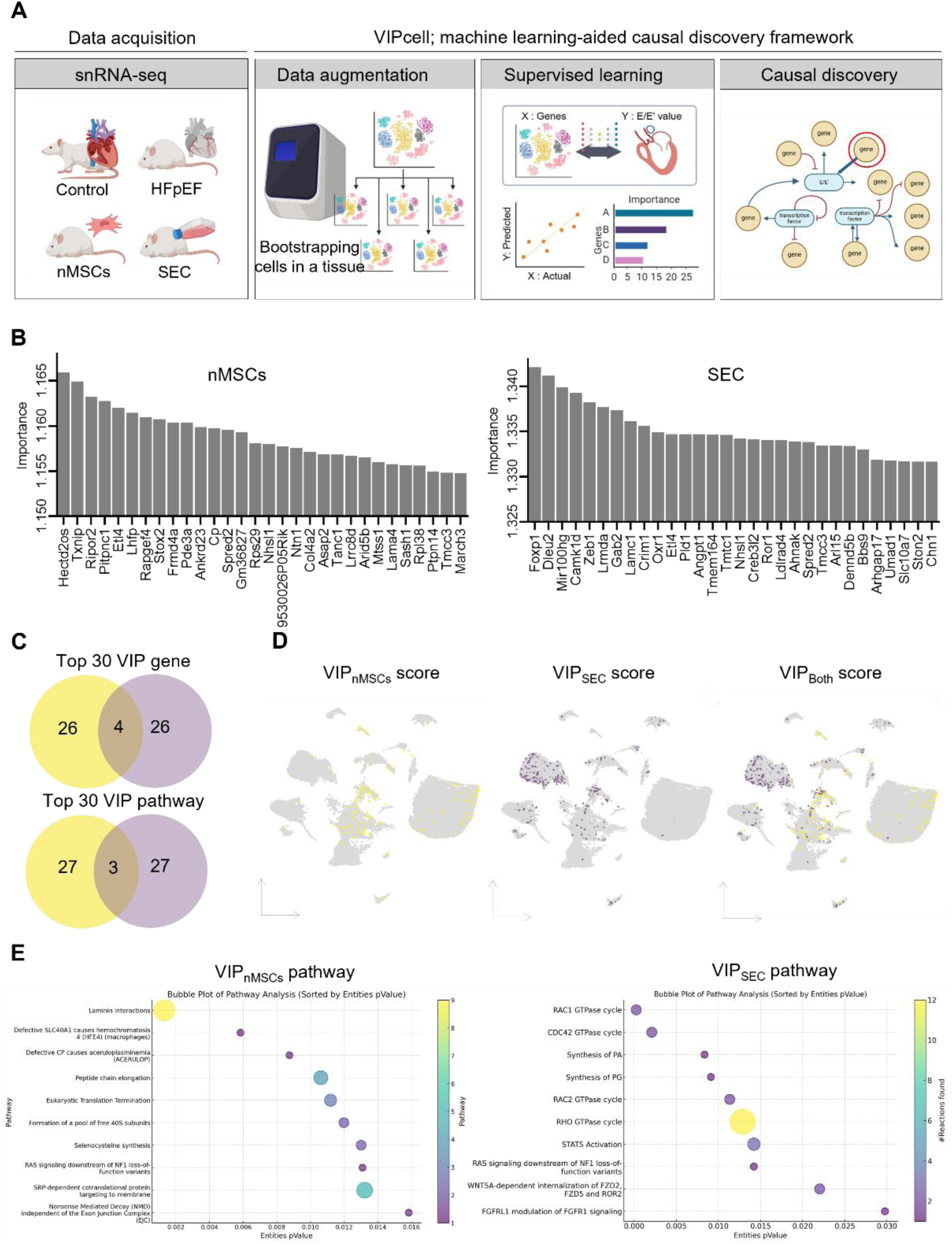
VIP genes identified by supervised learning on augmented snRNA-seq data of nMSCs and SEC groups. A: Schematic overview of VIP_cell_ framework. This novel approach integrates augmented bulkRNA-seq data from snRNA-seq data with tissue-level diastolic function measurements to identify key transcriptomic drivers and cell types associated with diastolic improvement in HFpEF. B: Bar graphs showing the calculated importances of the top 30 VIP genes identified by the VIPcell framework for observed E/E’ values in nMSCs and SEC-treated groups. The x-axis lists the genes, and the y-axis shows their importance scores. C: Venn diagrams showing the overlap of the top 30 VIP genes and their corresponding pathways between nMSCs and SEC groups. The upper Venn diagram shows the overlap of genes, while the lower Venn diagram shows the overlap of pathways. (Yellow; nMSCs, Purple; SEC). D: UMAP plots displaying the distribution of VIP_nMSCs_, VIP_SEC_, and VIP_shared_ genes expression across different cell clusters. The color intensity represents the VIP scores in each cell cluster. E: Bubble plots of pathway analysis for VIP_nMSCs_ and VIP_SEC_ genes. The size of the bubbles represents the number of entities (genes) involved in each pathway, and the color intensity represents the significance (p-value) of the pathway.

We employed a data augmentation approach as the first step of our framework to enhance the robustness of our machine learning model. By randomly bootstrapping four snRNA-seq datasets, we created 40,000 augmented datasets that maintained the original ground-truth transcriptomic information but were reassembled to simulate different tissue configurations. This approach generated diverse training datasets with preserved clustering patterns. We optimized cell sampling to ensure we captured meaningful variation and mitigate data redundancy. Unsupervised clustering showed that sampling 100 cells introduced excessive variability, clustering distinct groups like control and nMSCs together in principle component analysis. Conversely, sampling 10,000 cells resulted in minimal variation, undermining augmentation. Sampling 1,000 cells per tissue struck the optimal balance—retaining group-specific biological signals while introducing sufficient variability to enhance model generalizability(Supplement Fig. 9A). Model performance metrics validated this approach, with R² improving to 0.995 and MSE dropping to 0.07 as the number of augmented datasets increased to 20 (Supplement Fig. 9B, C). In contrast, the original four datasets alone failed to yield a predictive model, highlighting the critical role of augmentation in enabling ML-based modeling in low-sample single-cell studies.

As a second step, we applied PLS regression, a supervised learning technique that uncovers correlations between predictors and responses while providing a quantitative measure of predictor importance.^52–54^ This approach was used to establish two models for the nMSC and SEC groups. We prioritized the top 30 Variable Importance in Projection (VIP) genes contributing to E/e’ value at endpoint, out of 2,000 variable genes (Figure 4B). In agreement with relative nuclei fraction and cell-cell communication analyses, the top 30 VIP genes showed both shared and distinct patterns between the two groups. A Venn diagram analysis revealed that 4 out of the 30 VIP genes were common to both nMSCs and SECs, while the remaining genes were group-specific (Figure 4C). Further analysis of VIP gene expression distributions in 2D UMAP space showed that these genes localized in different clusters. Specifically, VIP_nMSCs_ genes were predominantly found in the EC I, CM I/II/III, Edc, LE, and Pc clusters, whereas VIP_SEC_ genes were mainly associated with the FB, Mc, Lym, EC I, Edc, Mu, SMC, SN, and CM IV clusters (Figure 4D). Notably, the EC I and Edc clusters were involved in heart remodeling post-HFpEF in both nMSC and SEC treatments, indicating shared remodeling contributions. Pathway enrichment analysis of the top 30 VIP genes using the Reactome Database further revealed both common and distinct regenerative mechanisms. In nMSCs group, pathways related to protein synthesis and laminin interaction were significantly enriched. In contrast, the SEC group showed enrichment in pathways linked to Rho GTPase signaling, immune modulation, and lipid metabolism, suggesting its potential to enhance remodeling by influencing cytoskeletal dynamics, the immune environment, and membrane biogenesis. These findings indicate that while nMSCs and their SEC share some remodeling pathways, they also engage distinct molecular mechanisms during heart failure recovery.

Causal discovery, a third step in the VIPcell framework, is an unbiased process used to infer causal structures between variables from observational data, without prior knowledge, to understand complex systems. It helps distinguish genuine causality from mere correlation, a focus of many machine learning methods like PLS regression.^55,56^ The DAS algorithm, a score-based method, evaluates the quality of causal graphs using a scoring function and effectively handles challenges such as hidden confounders, large datasets, and non-linear relationships common in biological systems, which traditional constraint-based causal discovery methods struggle with.^41,42^ We constructed causal structures between the top 30 VIP genes and the echocardiographic parameter E/e’ in both nMSCs and SEC groups using our curated dataset and the DAS algorithm (Figure 5). Each Directed Acyclic Graph (DAG) categorized genes as direct causes, indirect causes, or non-causal based on their relationship with E/e’. In the nMSCs graph, 10 out of the top 30 genes were identified as non-causal, including *Lhfp*, despite its high association rank. Among the 20 causal genes, *Pde3a*, *Txnip*, *Stox2*, and *Etl4* were identified as direct causes for E/e’. The DAG for the SEC group revealed five direct causes, seven indirect causes, and 18 non-causal genes. Notably, *Dleu2* and *Mir100hg*, highly associated with E/e’, were deemed non-causal, illustrating the limitations of conventional machine learning approaches that focus solely on association. The inference that five direct causal genes, *Foxp1*, *Arl14*, *Tmtc1*, *Camk1d*, and *Angpt1*, are initially induced by *Ston2*, a major component of the vesicle endocytosis machinery, is particularly noteworthy.^57^ Given that the secretome contains a large number of extracellular vesicles, the finding underscores the DAS algorithm’s reliability in identifying genuine causal relationships from the data. The functions of nine causal genes were investigated utilizing pathway enrichment analysis. Based on Reactome database, key enriched pathways include regulation of FOXO transcriptional activity, NLRP3 inflammasome, Tie2 signaling, NLR signaling pathways, purinergic signaling in leishmaniasis infection, and cell recruitment (pro-inflammatory response) (Supplement Fig. 10). These pathways collectively represent diverse biological processes involved in cellular signaling, immune regulation, and inflammatory responses. FOXO signaling is broadly involved in cellular homeostasis, differentiation, and metabolic adaptation, particularly its role in T cell maintenance, survival, and functional programming. FOXO transcription factors, particularly FOXO1 and FOXO3, regulate T cell quiescence, memory formation, and regulatory T cells (T_regs_) differentiation, influencing immune tolerance and responsiveness. ^58–61^ NLRP3 inflammasome activation and NLR signaling pathways are central to innate immune responses and can influence adaptive immunity through cytokine release and inflammasome-mediated signaling.^62,63^ Cell recruitment (pro-inflammatory response) plays a role in the mobilization of various immune cells to sites of inflammation. Among nine direct causal genes, *Foxp1*, *Arl15*, *Pde3a*, and *Txnip*, identified as the four causal genes closest to the E/e’ node, exhibit specificity in Cd4+ Foxp3+ T_reg_ cell activation. Foxp1, a key transcription factor, plays a crucial role in coordinating T_reg_ function by regulating Foxp3 chromatin binding, thereby maintaining T_reg_ cell identity and suppressive function. ^64,65^ Notably, Foxp1 is also involved in a regulatory feedback loop that modulates FOXO-driven transcriptional programs, which were significantly highlighted in our previous pathway enrichment analysis ^66^. *Arl15* encodes a small GTP-binding protein involved in TGF-β signaling, which has been positively correlated with Cd4+ T cell activity.^67–69^ *Txnip* is essential for T_regs_ stability and functions through the MondoA-Txnip axis, reinforcing immune homeostasis.^70,71^ *Pde3a*, a regulator of cyclic nucleotide signaling, modulates intracellular levels of cyclic adenosine monophosphate (cAMP) and cyclic guanosine monophosphate (cGMP), thereby contributing to Foxp3+ T-cell enrichment (Supplement Table 3). ^72,73^ The shared functional roles of these genes underscore the necessity for further investigation into the role of T_reg_ cells in HFpEF regeneration, particularly in the context of nMSC-based therapies and their secretome-derived factors. Moreover, these findings highlight the potential of causal machine learning in refining the selection of causally relevant genes from those highly associated in PLS regression, thereby enhancing mechanistic insights into T_reg_-mediated cardiac repair.

**Figure 5.**
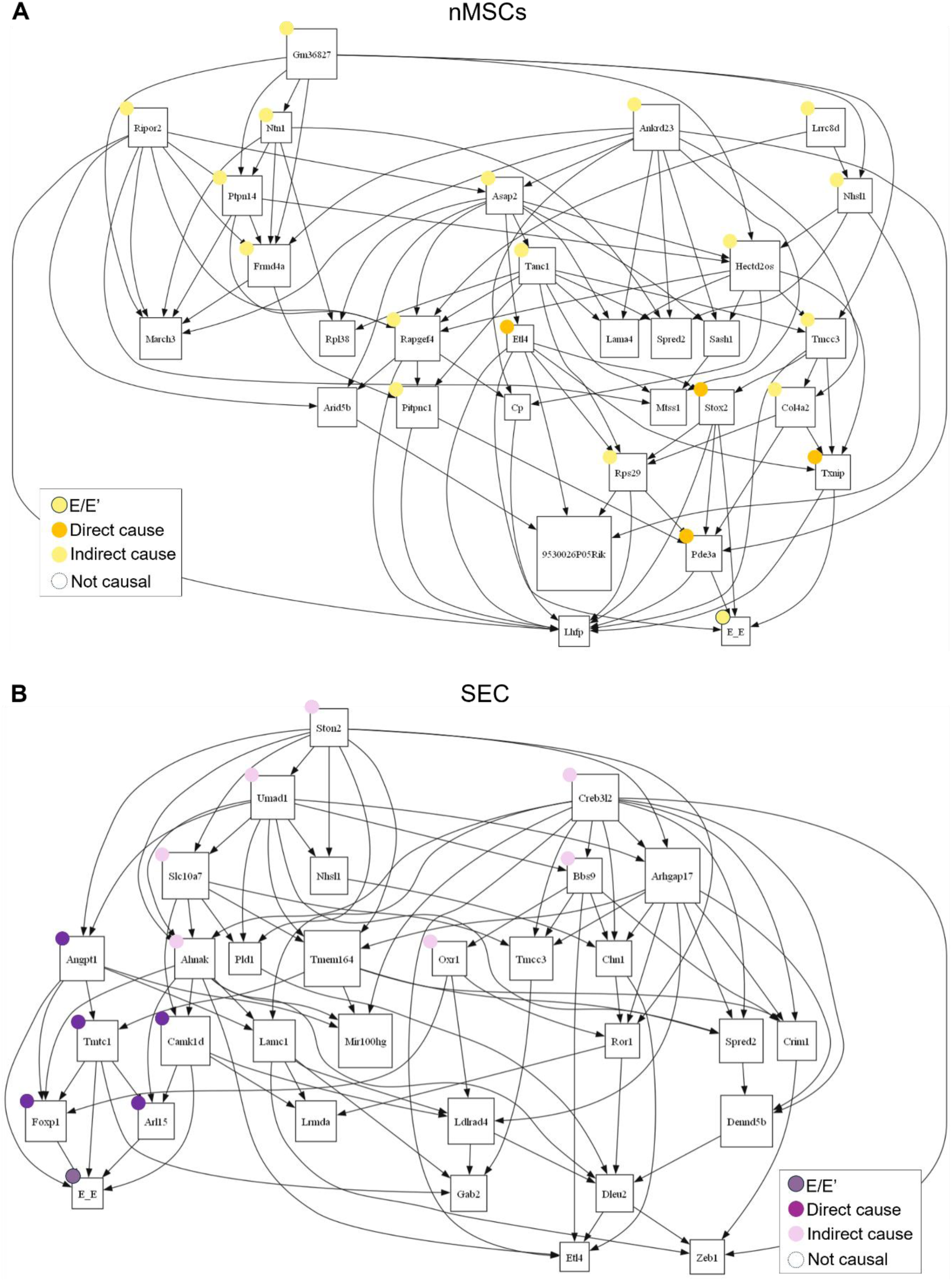
Advanced screening of causal VIP genes for diastolic function in nMSCs and SEC groups using a causal discovery algorithm. Directed acyclic graphs of top 30 VIP genes and E/E’ in A: nMSCs and B: SEC groups. Among top 30 VIP genes, identified causal genes are highlighted by dot on upper-left of gene labels. Edges means causes-and-effects relationship between paired genes and arrows direct from the source genes to the targets.

### Both nMSCs and SEC augment CD4^+^CD25^+^Foxp3^+^ T_regs_ in HFpEF and in vitro

To assess the immunomodulatory effects of nMSCs and SEC, we analyzed inflammatory and immune cell populations in the hearts of HFpEF HFD/L-NAME mice following treatment (Figure 6). T_reg_—specifically the CD4⁺CD25⁺FoxP3⁺ subset—are critical for immune homeostasis and suppression of inflammation. Natural T_reg_ are generated in the thymus via high-avidity interactions with MHC class II molecules, while adaptive (or induced) T_reg_ emerge in the periphery from naïve CD4⁺CD25⁻FoxP3⁻ precursors in response to appropriate stimulation. We first quantified CD4⁺CD25⁺FoxP3⁺ T_regs_ in mouse hearts. Compared to controls, HFpEF mice exhibited a significant reduction in cardiac T_regs_. Importantly, treatment with either nMSCs or SEC led to a significant increase in CD4⁺CD25⁺FoxP3⁺ T_regs_ relative to the untreated HFpEF group (Figure 6A,B), suggesting that both therapies promote a shift toward an anti-inflammatory immune profile. Next, we examined other immune cell populations in the myocardium, including total T cells (CD45⁺CD3⁺), helper T cells (CD45⁺CD4⁺), cytotoxic T cells (CD45⁺CD8⁺), B cells (CD45⁺CD19⁺), total macrophages (CD11b⁺CD64⁺F4/80⁺), tissue-reparative macrophages (CD11b⁺CD64⁺F4/80⁺CD206⁺), and tissue-resident macrophages (CD11b⁺CD64⁺F4/80⁺TIMD4⁺). No significant differences in these populations were observed across the control, HFpEF, nMSC, or SEC groups (Figures 6B-F, Supplement Fig. 11).

**Figure 6.**
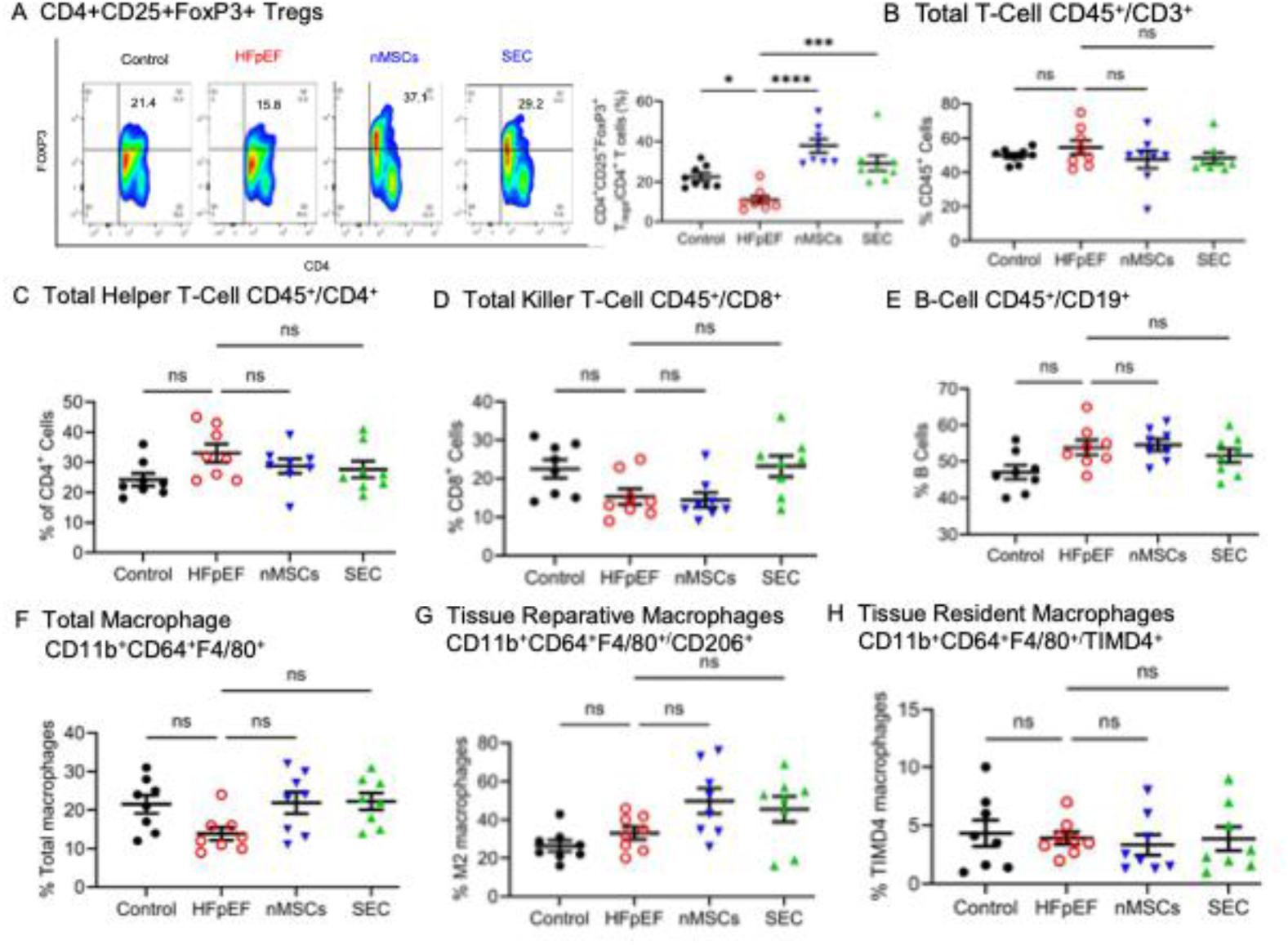
Both nMSCs and SEC augmented cardiac CD4^+^CD25^+^FoxP3 T_reg_ cell subsets in the hearts of mice of HFpEF and control and nMSC stimulate T_regs_ in vitro. A and B: Flow cytometric analysis of cardiac CD4^+^CD25^+^FoxP3^+^ regulatory T cells (FoxP3 T_reg_) in the mice of control, HFpEF, nMSCs, and SEC groups (n = 5-6 hearts/group). Flow cytometric analysis of C: Total T-Cell CD45^+^/CD3^+^; D: Total Helper T-Cell CD45^+^/CD4^+^; E: Total Killer T-Cell CD45^+^/CD8^+^; F: B-Cell CD45^+^/CD19^+^ (n = 5-6 hearts/group). G and H. Flow cytometric analysis of cardiac CD4^+^CD25^+^FoxP3^+^ regulatory T cells (FoxP3 T_reg_) when nMSCs, CDCs, aCPCs, or BM-MSCs are co-cultured with human T cells (n = 5-6 hearts/group). Data was analyzed by Ordinary one-way ANOVA using Prism software (V 10). Holm-Sidak’s test was used for multiple comparisons. * p<0.05, **p<0.01, ns = not significant

Together, these findings suggest that nMSCs and SEC selectively expand the T_regs_ population in the hearts of HFpEF mice, potentially contributing to their anti-inflammatory and cardioprotective effects. To determine whether the expansion of T_regs_ is specific to nMSCs, an in vitro head-to-head comparison was performed using human T cells co-cultured with nMSCs or other well-characterized stem/progenitor cell types, including adult bone marrow–derived mesenchymal stromal cells (BM-MSCs), adult cardiosphere-derived cells (CDCs), adult cardiac progenitor cells (aCPCs), or a placebo control containing cell-free Iscove’s Modified Dulbecco’s Medium (IMDM) (Figures 6G and H). After 4 days of co-culture, flow cytometry analysis revealed that nMSCs significantly increased T_reg_ levels compared to aCPCs, BM-MSCs, or the control. Notably, CDCs also induced a significant increase in T_regs_, although to a significantly lesser extent than nMSCs. These results demonstrate that nMSCs possess a unique and superior capacity to stimulate T_reg_ expansion compared to other adult stem/progenitor cells.

### A decrease in cardiac CD4^+^CD25^+^FoxP3^+^ T_reg_ blocks the cardioprotective effects of both nMSCs and SEC on HFpEF mice

To determine whether cardiac CD4⁺CD25⁺FoxP3⁺ T_reg_ mediate the therapeutic effects of nMSC and SEC treatment in HFpEF, we investigated the impact of T_reg_ depletion on myocardial fibrosis and cardiac function. We utilized FTY720 (fingolimod), an immunomodulatory agent and sphingosine-1-phosphate receptor (S1PR) modulator, which induces internalization of S1P1 receptors upon phosphorylation by sphingosine kinase, effectively sequestering lymphocytes in lymph nodes and preventing their peripheral circulation.^74,75^ As outlined in Figure 7A, mice in the control-FTY720, HFpEF-FTY720, nMSC-FTY720, and SEC-FTY720 groups received daily intraperitoneal injections of FTY720 (3 mg/kg) for 18 consecutive days, starting 10 days after the first dose of nMSCs, SEC, or vehicle injection. Control animals received equal volumes of normal saline (N.S.). Flow cytometry revealed that treatment with FTY720 significantly reduced cardiac CD4⁺CD25⁺FoxP3⁺ T_regs_ across all FTY720-treated groups—including control, HFpEF, nMSCs, and SEC—compared to their respective N.S.-treated counterparts (Figure 7B). We next assessed myocardial fibrosis using Masson’s trichrome staining. As expected, fibrosis was significantly elevated in the HFpEF-N.S., HFpEF-FTY720, nMSC-N.S., nMSC-FTY720, SEC-N.S., and SEC-FTY720 groups compared to the control-N.S. group. However, there were no significant differences in fibrosis between control-N.S. and control-FTY720, HFpEF-N.S. and HFpEF-FTY720, nMSC-N.S. and nMSC-FTY720, or SEC-N.S. and SEC-FTY720 groups (Figure 7C), indicating that T_reg_ depletion alone did not exacerbate fibrosis beyond that already induced by HFpEF. We then evaluated diastolic function via mitral E/e′ ratio measurements. While no significant differences were observed between control-N.S. and control-FTY720 or HFpEF-N.S. and HFpEF-FTY720, both the nMSC-FTY720 and SEC-FTY720 groups exhibited a significantly higher E/e′ ratio compared to their respective nMSC-N.S. and SEC-N.S. groups (Figure 7D).Taken together, these data indicate that depletion of cardiac CD4⁺CD25⁺FoxP3⁺ T_reg_ using FTY720 abrogates the beneficial effects of nMSC and SEC therapy on myocardial fibrosis and diastolic function in HFpEF mice. These findings highlight T_reg_ as a key mechanistic link in the cardioprotective effects of stem cell-based therapies in HFpEF.

**Figure 7.**
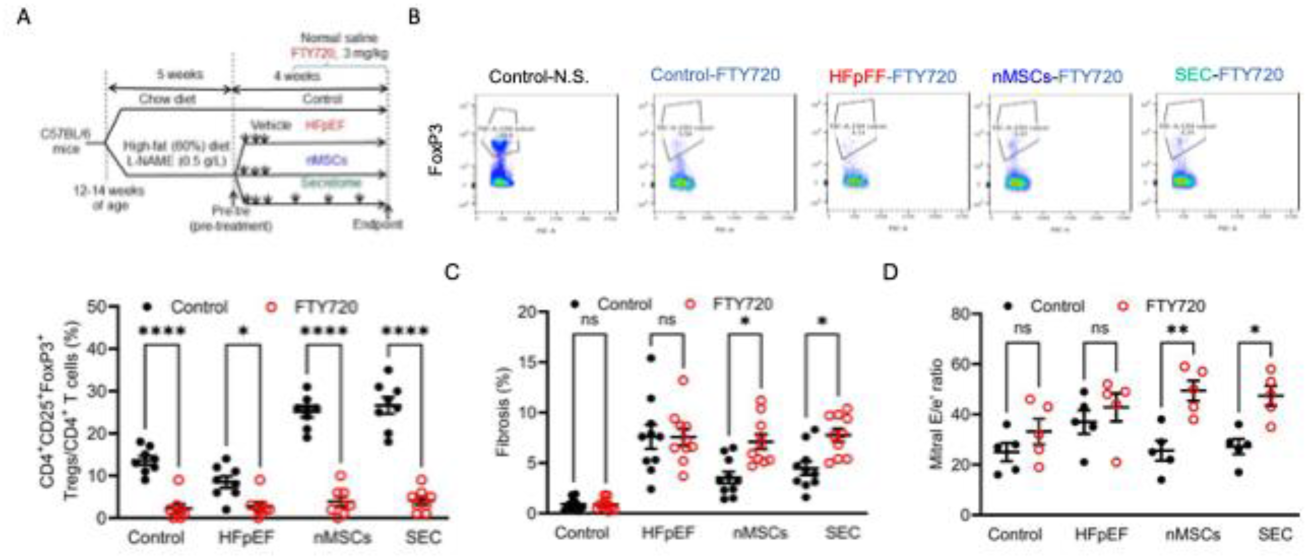
Chemical depletion of CD4^+^CD25^+^FoxP3^+^ T_reg_ blocked the protective effects of nMSCs and secretome in HFpEF. A: Experimental procedures. The mice of control, HFpEF, nMSCs, and secretome (SEC) groups were treated with normal saline (N.S.) or FTY720 for 18 days. B: flow cytometric analysis of cardiac CD4^+^CD25^+^FoxP3^+^ T_regs_ in the mice of control, HFpEF, nMSCs, and SEC groups treated with N.S. or FTY720 (n = 8 mice/group). C: quantification of myocardial fibrosis in the mice of control, HFpEF, nMSCs, and SEC groups treated with N.S. or FTY720 (n = 10 heart sections/group). D: quantification of mitral e/e’ ratio in the mice of control, HFpEF, nMSCs, and SEC groups treated with N.S. or FTY720 (n = 5 mice/group). Data was analyzed by Ordinary one-way ANOVA using Prism software (V 10). Holm-Sidak’s test was used for multiple comparisons. * p<0.05

## Discussion

In this study, we demonstrate that both nMSCs and the SEC significantly improve cardiac diastolic function, reduce myocardial fibrosis, and enhance exercise capacity in two distinct murine models of HFpEF. These therapeutic benefits were linked to a shift in the myocardial immune landscape toward an anti-inflammatory state and were mechanistically dependent on CD4^+^CD25^+^FoxP3^+^ T_reg_. Through integrative snRNA-seq and a novel causal machine learning framework (VIPcell), we uncovered a unifying immunoregulatory axis driving cardiac recovery.

This has significant therapeutic implications; especially as chronic inflammation is increasingly recognized as a key driver of disease progression and poor outcomes in HFpEF.^76^ Despite this, previous efforts to reduce inflammation using broad-spectrum immunosuppressive agents have largely fallen short in clinical trials. Against this backdrop, our results point to a novel and more targeted mechanism of action: recovery mediated by T_reg_, which play a central role in suppressing inflammation and limiting fibrosis. Importantly, the observed improvements in early diastolic relaxation (e′ velocity) and exercise capacity suggest that nMSC and SEC therapies enhance intrinsic myocardial relaxation—a therapeutic domain not addressed by current HFpEF treatments. Existing drug classes, including ARNI, SGLT2 inhibitors, GLP-1 receptor agonists, and MRAs, primarily act via decongestion. Our findings introduce a new avenue for therapeutic development aimed at directly restoring myocardial function and chronic inflammation.

Although both nMSC and SEC therapy improved diastolic performance, they exerted distinct remodeling signatures at the cellular level. nMSC treatment broadly restored cardiac cell type composition toward a homeostatic normal state, particularly rebalancing endothelial and immune populations. In contrast, SEC therapy preferentially expanded cardiomyocyte subpopulations and activated more focused signaling between endothelial cells and cardiomyocytes. These divergent remodeling patterns underscore the versatility of nMSC-derived interventions, where intact cells and their secreted products may engage the heart via overlapping but non-identical pathways. A common mechanistic denominator, however, emerged from both treatments: selective expansion of CD4⁺CD25⁺Foxp3⁺ T_reg_. Flow cytometry confirmed increased T_reg_ abundance in treated hearts, suggesting that both interventions shift the immune balance toward regulatory rather than effector phenotypes. Notably, head-to-head in vitro assays revealed that nMSCs uniquely and potently induced T_reg_ expansion compared to four other adult stem/progenitor cell types, including BM-MSCs and CDCs—highlighting a distinct immunoregulatory capacity intrinsic to neonatal-derived cells. These findings align with our prior work demonstrating intravenous delivery of nMSCs can increase resident T_reg_ within the myocardium in a model of HFrEF.^22^ We have further determined that nMSC exosomal delivery of a specific microRNA was alone sufficient to promote T_reg_ activation and enhance their immunoregulatory function (Data not shown). Collectively, these findings highlight T_regs_ as a nMSCs-specific central effector population and a promising target for immunomodulatory therapy across heart failure phenotypes.

To systematically elucidate the molecular underpinnings of these effects, we developed VIPcell, a machine learning framework that integrates data augmentation, supervised learning, and scoring-based causal inference. By generating biologically meaningful augmented samples from limited, high-dimensional single-cell data, VIPcell bridges gene expression, cellular context, and organ-level phenotypes. Using this framework, we identified gene regulators not only associated with, but likely causal for, improved diastolic function as measured by E/e′. Among these, Foxp1, Txnip, Pde3a, and Arl15—all known to influence T_reg_ biology—were validated through causal modeling and experimental confirmation.

Compared to traditional approaches such as transcriptome-wide association study^77,78^ or gene regulatory network inference^79^, VIPcell adopts a multiscale, directionally resolved strategy for mechanistic gene prioritization. By integrating Partial Least Squares (PLS) regression with the Discovery At Scale (DAS) algorithm, VIPcell screens important genes for a specific function and infers causal graphs that capture directional gene–phenotype relationships, even under high-dimensional and nonlinear conditions. DAS offers advantages over constraint-based or SEM methods by addressing hidden confounding and enhancing scalability.^41,42^ Unlike single-cell GRN models like CellOracle^80^ or LINGER^81^, which remain confined to cell-intrinsic or cell fate dynamics, VIPcell infers causal graphs through bootstrapped resampling and probabilistic scoring—directly linking transcriptomic signals to ventricular function. The resulting identification of direct gene–function axes, such as the Foxp1(gene)–T_reg_(cell)–E/e’ (diastolic function) link, highlights the translational utility of this platform. As recent causal frameworks in regenerative immunology and developmental systems demonstrate, such mechanistic insight is critical for precision targeting of gene- or cell-based therapies.

This work expands on prior regenerative therapy studies in HFpEF in several keyways. Previous investigations have shown that various progenitor cell types can confer anti-fibrotic or functional benefits in animal models, but most lacked mechanistic clarity and did not resolve the identity of key cellular effectors.^82,83^ Here, we identify T_reg_ as central to recovery and define their causal gene networks using a framework specifically adapted for high-dimensional single-cell data. Our approach also reveals that secretome therapy—an acellular alternative—can recapitulate many benefits of live cell transplantation. This finding supports the feasibility of scalable, off-the-shelf biologics for complex syndromes like HFpEF, where logistical challenges limit the utility of other cell therapies.

From a translational perspective, these results highlight several clinical implications. First, our data suggest that measuring T_reg_ levels or activity may serve as a pharmacodynamic biomarker in cell-based or secretome-based trials. Second, the identification of causal T_reg_-associated genes provides candidate targets for adjunctive therapies, such as low-dose IL-2 or small molecules that stabilize T_reg_.^84,85^ Third, given the conserved nature of MSC-derived vesicles and cytokines, SEC-based products may enable immune-tolerant, cross-species application, accelerating therapeutic development.^18^

This study has limitations. First, HFpEF is a multifactorial syndrome involving aging, systemic inflammation, metabolic dysfunction, and cardiovascular comorbidities. While the combination of a HFD and L-NAME reliably induces key HFpEF features, such as diastolic dysfunction and systemic inflammation, neither this model nor the HFpEF db/db model adequately recapitulates aging—a major contributor to HFpEF pathophysiology in humans—which may limit generalizability. Second, our follow-up period was relatively short. Given the robust improvements observed within one month of treatment, we selected an early endpoint to minimize potential confounding from secondary remodeling effects. However, this approach precludes assessment of the long-term durability or safety of either treatment, which should be addressed in future studies. Third, our single-nucleus dataset, while informative, may miss dynamic or post-transcriptional effects not captured at the transcript level. VIPcell remains to be tested across broader disease models. Last, we did not isolate the active components within the secretome; future studies dissecting exosomes and independently secreted protein fractions will be essential to identify the most therapeutically relevant cargo.

In summary, this study provides mechanistic evidence that nMSC and SEC therapies improve HFpEF through immune reprogramming and T_reg_-mediated cardiac repair. By integrating single-nucleus transcriptomics with causal machine learning and in vivo validation, we identify a conserved immunoregulatory axis as the common therapeutic denominator across both interventions. These insights support the development of data-driven, T_reg_-centered therapies and position nMSC secretome as a promising, scalable approach for immunomodulatory treatment in HFpEF.

## METHODS

### Sex as a biological variable

Only male mice were used in the experiments as female mice are less susceptible for the treatment of high-fat-diet (HFD) and L-N^G^-nitroarginine methyl ester (L-NAME).^24^ In the db/db mouse study, our study examined male and female animals, and similar findings are reported for both sexes.

### Induction of HFpEF in mice

Male C57BL/6 mice were provided by The Jackson Laboratory (Bar Harbor, ME). They were cared in the facility of Northwestern University Chicago Campus (Chicago, IL). The animals were kept on a 12-h light-dark cycle in a temperature-controlled room. Only male mice were used in the experiments as female mice are less susceptible for the treatment of high-fat-diet (HFD) and L-N^G^-nitroarginine methyl ester (L-NAME).^24^ The experimental procedures were approved by the Animal Care and Use Committee of Northwestern University and conformed to the National Institutes of Health Guide for the care and use of Laboratory Animals (NIH Publications No. 8023, 8th edition, 2011).

The murine model of HFpEF was established, as described.^25^ Briefly, male C57BL/6 mice at 12-14 weeks of age were induced to develop HFpEF by feeding a HFD where 60% of the calories were from fat (lard and soybean oil), 20% from proteins, and 20% from carbohydrates (D12492, Research Diets, Inc) and water spiked with L-NAME (0.5 g/L; Sigma Aldrich, Catalog #: N5751-10G). Our preliminary study indicated that 75% of the mice developed HFpEF at the time point of 5 weeks after feeding HFD and L-NAME. Since 80% of the mice would not remain HFpEF 4 weeks after the withdrawal of HFD and L-NAME, the mice were fed both HFD and L-NAME throughout the study.

In addition to above cardiometabolic HFpEF, we used another murine model driving by type 2 diabetes, obesity, hyperlipidemia, and pulmonary congestion, since a significant cohort of HFpEF patients includes aging, obese individuals with type 2 diabetes, hyperlipidemia, pulmonary congestion, and skeletal muscle weakness.^26–28^ Lepr db/db mice (db/db mice), a leptin receptor-deficient model of obesity and type 2 diabetes, and their nondiabetic lean heterozygous Lepr db/+ littermates (db/+ mice) were purchased from The Jackson Laboratory. An equal number of male and female mice were used and fed normal chow throughout the experiments.

### Experimental protocols

To determine the therapeutic effects of nMSCs and SEC on HFpEF, C57BL/6 mice were randomly assigned into the following 4 groups: control, HFpEF, HFpEF+nMSCs, and HFpEF+SEC (Figure 1A). In the control group (n = 14 mice), mice at 12-14 weeks of age had unrestricted access to a standard murine chow diet of 19.1% protein, 6.5% fat, and 47% carbohydrates (2020X from Teklad) and water throughout 9 weeks of the study. The other 3 groups of mice (n = 50 mice) had unrestricted access to HFD and water containing 0.5g/L of L-NAME throughout the study. After establishing the HFpEF model at 5 weeks with HFD and L-NAME (Supplement Fig.1), we optimized the dosing of the nMSCs and SEC by increasing the dosage of nMSCs from 10 M/kg to 50 M/kg and SEC from 1mg/kg, 10 mg/kg, and then 50 mg/kg (Supplement Fig. 3) and subsequently arrived at the optimal dosing of nMSCs at 50 M/kg and SEC at 50 mg/kg with maximal recovery recorded. After developing HFpEF (n = 38 mice) 5 weeks after HFD and L-NAME, the mice were injected phosphate-buffered saline (PBS, n=14) HFpEF control group, 50 × 10^6^/kg nMSC group (n=14), or 50 mg/kg proteins for the SEC group (n=14) through tail vein. PBS, nMSCs, and SEC were injected 1 time every 48 hours for 6 days and subsequently 1 time every 7 days for the next 3 weeks. Body weight of each mouse was measured weekly for 9 weeks at the same time of day. Blood glucose was measured with an AimStrip plus blood glucose meter (Germaine Laboratories, Inc., San Antonio, TX) after mice were fasted for 12 hours. The left ventricle of mice was evaluated using an echocardiography at the time points of baseline (before feeding HFD and L-NAME), fed HFD and L-NAME for five weeks, and then treatment with PBS, nMSCs, or SEC, and then thereafter at 1, 2, and 4 weeks after the treatment of PBS, nMSCs, and SEC, as described below. HFD and L-NAME were fed throughout the study. Similar experiments were performed in the db/db and db/+ mice using placebo, nMSCs, or SEC at 48 weeks of age when diastolic dysfunction was present and then extended for another 4 weeks with the treatment groups (Supplement Fig 2-3).

To determine the contribution of CD4^+^CD25^+^FoxP3^+^ T_reg_ to the cardioprotective effect of nMSCs and SEC in HFpEF, we used FTY720 (Sigma-Aldrich), a known depletory agent of CD4^+^CD25^+^FoxP3^+^ T_reg_^29^ to deplete CD4^+^CD25^+^FoxP3^+^ T_reg_ in a separate cohort of the 4 groups of mice: control, HFpEF, nMSCs, and SEC (Figure 7A). All mice were intraperitoneally injected 3 mg/kg FTY720 daily for 18 days, starting at 10 days after the treatment of PBS, nMSCs, or SEC. The left ventricle was evaluated with an echocardiography, myocardial fibrosis was quantified in Masson’s chrome-stained heart sections, and CD4^+^ T cells and CD4^+^CD25^+^FoxP3^+^ T_regs_ were quantified on a flow cytometers.

### Transthoracic echocardiography

Non-invasive transthoracic echocardiography was used to evaluate the left ventricle in C57BL/6 mice with and without HFpEF by a reader blinded to HFpEF and treatment status. Animals were sedated by the inhalation of 1.5 % isoflurane and oxygen. Echocardiography was performed with a VisualSonics Vevo 3100 High-resolution Imaging System (Toronto, Canada), as we have previously described.^30^ Left ventricular dimensions was measured by two-dimension guided M-mode method. Left ventricular ejection fraction was calculated from the following equation: (end-diastolic volume – end-systolic volume)/end-diastolic volume × 100. Pulsed Doppler waveforms recorded in the apical 4-chamber view were used for the measurement of the peak velocities of mitral E (early diastolic mitral inflow) wave. Tissue Doppler waveforms recorded in the apical 4-chamber view were used to measure mitral é (early diastolic mitral annulus) velocity. Mitral E/ é ratio was used for the evaluation of LV diastolic function as a marker of LV filling pressure.

### Treadmill exercise stress test

Mice ran on a Panlab 76-1183 Two-Lane Treadmill (Harvard Bioscience, Inc.) with Stimulus Detection on a 20° upward incline. The day before the test, animals were subjected to an acclimation at 2.4 m/min for 3 min. Mice were run for 5 min at a speed of 5 m/min, then for 2 min at 14 m/min. The speed was increased by 2 m/min every 2 min until mice reached exhaustion. Exhaustion is defined as the animal getting shocked continuously for 5 s. The Treadmill Software measured running distance, running duration, and exhaustion.

### Histopathological examination of mouse hearts

To validate the findings from echocardiographic examination of mice, mice at 4 weeks after the treatment of nMSCs or SEC were euthanized, and mouse hearts were visualized and weighed. Dehydrated mouse hearts were embedded with paraffin and sliced transversely from the apex to the basal part of the left ventricle at 4-5 μm-thickness. Sections were stained with Masson’s trichrome to assess myocardial fibrosis and wheat germ agglutinin (WGA) for cardiomyocyte surface area (n = 10 sections/mouse, 3 mice/group).^31^ Stained sections were imaged with an Olympus microscope (Olympus America, Melville, New York) at 200× magnification or as indicated. Image analysis was completed with ImageJ software.

For Masson’s Trichrome staining, isoflurane-anesthetized mice received intracardiac saturated KCl (30 mmol/L and 5% dextrose in 1× PBS) to arrest the heart in diastole, as described. Mouse hearts were washed with cold PBS, fixed with 10% Formalin, dehydrated, embedded in paraffin, and stained with Masson’s trichrome. The percentage of total fibrosis area was calculated as the summed, blue-stained regions of interest divided by total area. For immunohistochemical staining, cardiomyocyte size was measured by staining mouse hearts with WGA. Mouse hearts were fixed with 4% paraformaldehyde then embedded in paraffin. Sections from the midpoint of the left ventricle were deparaffinized and hydrated. Tissue sections were stained with fluorescence Oregon Green 488–labeled WGA.

### Flow cytometry

Mouse hearts were flushed with PBS, and the left ventricle was then excised, minced, and digested with collagenase D (Roche) at 37 °C for 50 minutes on a rocking platform (180–200 rpm). After enzymatic digestion, the cell suspension was filtered through a 70-μm cell strainer (Fisher Scientific #22363548) and centrifuged at 500×g for 10 minutes. To lyse red blood cells, the cell pellet was incubated in ammonium-chloride-potassium lysing buffer (Gibco # A10492–01) at room temperature for 3–5 minutes, then washed with iced cold fluorescence activated cell sorter washing buffer (2.5% fetal bovine serum in PBS without calcium and magnesium). Cells were resuspended in the washing buffer, and samples were incubated with Fc-Block (anti-rat CD16/CD32, 0.5 μg per 1 million cells) before incubation with isotype controls or primary antibodies, according to the manufacturer’s instructions. Cells were then washed with washing buffer. Flow cytometry was performed on a BD-LSRFortessa cytometer (BD Biosciences, San Jose, CA) and data were analyzed by FlowJo software (Tree Star, Inc., Ashland, OR), as described.^32^ The antibodies are described in Supplement Table 1. T cells and T_reg_ were first gated (FSC-A vs SSC-A) as lymphocytes. For total T cells, the CD3 cells were gated and further analyzed for CD4 and CD8. For T_reg_, CD4 cells were gated, from this gate, CD25^+^ and FoxP3^+^ double-positive cells were determined.

### snRNA-seq analysis

Randomized four mice from each group of control, HFpEF, nMSCs, and SEC were confirmed to have an appropriate change in diastolic function confirmed by echocardiogram in the HFD/L-NAME model of HFpEF and were euthanized with CO_2_, and the hearts were rapidly trimmed of large vessels connected to the hears on ice. Nuclei were extracted from cardiac tissues. as described.^33^ Isolated nuclei were enriched as per instructions from 10x Genomics “Nuclei Isolation from Cell Suspensions & Tissues for Single Cell RNA Sequencing” protocol (CG000124, Rev F) to reach the target concentration of 700–1200 nuclei/µl. The suspension was processed using 10X Genomics Single Cell protocol (CG00053, Rev D).

### snRNA-seq Processing

Nuclei number and cell viability were analyzed using Nexcelom Cellometer Auto2000 with AOPI fluorescent staining method. Sixteen thousand nuclei were loaded into the Chromium iX Controller (10X Genomics, PN-1000328) on a Chromium Next GEM Chip G (10X Genomics, PN-1000120), and processed to generate single-cell gel beads in the emulsion (GEM) according to the manufacturer’s protocol. cDNA libraries were generated using the Chromium Next GEM Single Cell 3’ Reagent Kits v3.1 (10X Genomics, PN-1000286) and Dual Index Kit TT Set A (10X Genomics, PN-1000215) according to the manufacturer’s manual. Quality control for constructed libraries were performed with the Agilent Bioanalyzer High Sensitivity DNA kit (Agilent Technologies, 5067-4626) and Qubit DNA HS assay kit for qualitative and quantitative analysis, respectively. The multiplexed libraries were pooled and sequenced on Illumina Novaseq6000 sequencer with 100 cycle kits using the following read length: 28 bp Read1 for cell barcode and UMI, and 90 bp Read2 for transcript. The single cell library preparation and sequencing was done at Northwestern University NUseq facility core with the support of NIH Grant (1S10OD025120). Raw sequencing data, in base call format (.bcl) was demultiplexed using Cell Ranger from 10x Genomics, converting the raw data into FASTQ format. Cell Ranger was also used for count quantification and alignment to the mouse reference genome (mm10). The resulting matrix files which summarize the alignment results were imported to Seurat v.4 (Satija Lab, NYGC) for further analysis.^34^

### snRNA-seq Data Analysis

Seurat objects were created from the Cell Ranger output using a per cell minimum feature setting of 100. Cells with less than 200 UMIs, less than 300 unique genes, and having a mitochondrial ratio > 0.05 were also excluded from downstream analysis. Read counts were normalized using the SCTransform function, with mitochondrial ratio regressed out during the process. Datasets for all four groups were integrated using default parameters. A principal component (PC) analysis was performed on the integrated dataset to identify highly variable genes. The top 30 PCs were then used for clustering projection using a uniform manifold approximation and projection algorithm (UMAP). Clustering analysis was performed with a resolution parameter of 0.4. Markers for each cluster were found using the FindAllMarkers function with a logFC threshold of 0.25 and a return threshold of 0.05. Clusters were identified by cross-referencing their highest expressed markers to the CellMarker (v 2.0)^35^, PanglaoDB (https://panglaodb.se/)^36^, and CZ CELLxGENE (https://cellxgene.cziscience.com/)^37^ databases. When necessary, we also searched markers through a manual literature search. For heatmaps, the integrated, scaled, and normalized dataset was down sampled to 100 cells and the Seurat package’s DoHeatmap function was used to plot the differentially expressed genes in each cluster.

### Cell-cell Interaction Inference

The R package, CellChat (v 2.0)^38^ was utilized to infer global cell-cell communication patterns. Following the given vignette from CellChat (http://www.cellchat.org/), Seurat objects from the nMSCs and SEC snRNA-seq data were transformed into CellChat objects. CellChatDB, database encompassing 2,021 validated molecular interactions, was used for inference. Communication pathway probabilities were calculated using ‘computeCommunProb-Pathway’ with significance thresholds set at p < 0.05. To visualize global patterns and probability heatmap between identified clusters, built-in plotting functions are employed, following the provided tutorials.

### snRNA-seq Data Augmentation and Conversion

A data augmentation strategy was implemented to expand single nucleus RNA-sequencing (snRNA-seq) dataset using Jupyter Lab and Python (v 3.11). The original dataset comprised four snRNA-seq samples, collectively containing approximately 26,410 nuclei. In the augmentation process, 1,000 cells from each sample’s curated matrix were randomly selected, along with their corresponding gene expression profiles. The normalized gene expression values of these selected cells were then averaged to generate augmented bulk RNA-seq data. This procedure was iteratively repeated 10,000 times per group to create a comprehensive collection of augmented data, which was subsequently compiled and exported as a comma-separated values (.csv) file for further computational modeling analysis.

### Partial Least Square Regression

For computational modeling, the augmented bulk RNA-seq dataset was integrated with each group’s averaged E/e’ values from echocardiograms for PLS regression. NumPy and Pandas for data manipulation and structuring, Matplotlib for data visualization, and various modules from Scikit-learn for machine learning tasks were employed in python. The PLS regression model was implemented using Scikit-learn’s PLSRegression class (components number = 10), with model performance evaluated using cross-validation techniques (RepeatedKFold, Fold = 5) and metrics such as mean squared error and R^2^ score. Each gene’s variable importance in projection (vip) scores were calculated from this model. vip scores were derived from the model’s scores (*t*), weights (*w*), and loadings (*q*). To compute the importance score for each feature, the following formula was used.

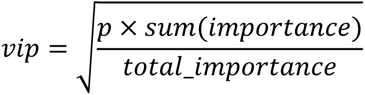

where *importance* is the diagonal of the matrix product involving (*t*), (*q*), and (*q^T^*), and (*p*) is the number of features. Two distinct PLS regression models were developed using augmented 30,000 RNA-seq data points from control, HFpEF, and nMSC or SEC snRNA-seq datasets, focusing on 2,000 selected genes linked to functional outcomes. The top 30 genes of nMSCs and SEC group were ranked by their vip score and plotted on the clustering projection by ‘feature plot’ function in R package, Seurat. These genes were then subjected to pathway enrichment analysis using the Reactome^39,40^ web-based tool (https://reactome.org/). The top 30 pathways were identified and ranked based on their statistical significance, as determined by p-value.

### Scoring-based Causal Discovery

Causal discovery was sequentially performed to uncover the cause-and-effect structure among the top 30 VIP genes and the E/e’ value at endpoint using 30,000 augmented data points. Utilizing the ‘dodiscover (v 0.0)’ library, a context for causal discovery was created and applied the DAS algorithm^41,42^ to infer the causal structure between these genes and the functional outcome. In this framework, each variable is modeled as part of an additive structural causal model, where the goal is to infer a directed acyclic graph (DAG) that captures the underlying causal structure. The DAS algorithm first estimates a topological ordering of variables using the SCORE approach, then sequentially tests and assigns directed edges based on statistical criteria, followed by a pruning step to reduce false positives. The resulting graphs were visualized using the ‘pywhy_graphs.viz’ module. In these graphs, genes were categorized as ‘direct cause’, ‘indirect cause’, or ‘not causal’ based on their linking edges to the E/e’ node. Among the top 30 genes, only those identified as direct causes were annotated as VIP causal genes.

## Acknowledgements

SK is funded through NIH 1R01HL118491, and Lurie Children’s Hospital fund. MED and JH are funded by the Additional Ventures Cures Collaborative.

