## Supplemental Figures for "Machine Learning Identifies Distinct Treg-Mediated Remodeling in HFpEF Hearts Treated with Neonatal Mesenchymal Stem Cells and Their Secretome"

**Table of Contents**

1. **Supplement Figure 1.** Induction of HFpEF by HFD and L-NAME in C57BL/6 mice.
2. **Supplement Figure 2.** Optimal dosing of nMSCs or their SEC for treating HFpEF.
3. **Supplement Figure 3.** Schematic Study Design in db/db mice.
4. **Supplement Figure 4.** Induction of HFpEF in db/db mice.
5. **Supplement Figure 5.** Therapeutic efficacy in diastolic function of nMSCs or SEC in HFpEF db/db and db/+ mice.
6. **Supplement Figure 6.** Cardiac functional assessment in db/db and db/+ mice.
7. **Supplement Figure 7.** Effect of nMSC and SEC on exercise tolerance of db/db induced HFpEF mouse model.
8. **Supplement Figure 8.** Both nMSCs and SEC did not significantly change increased cardiomyocyte size in HFpEF mice.
9. **Supplement Figure 9**. VIPcell framework enabled supervised and causal machine learning on limited number of snRNAseq data for measured tissue-level diastolic function.
10. **Supplement Figure 10.** Pathway enrichment analysis of nine direct causal genes for E/e' diastolic function identified from nMSCs- and SEC-treated HFpEF groups.
11. **Supplement Figure 11.** Effect of nMSCs- or SEC-treatment on various types of macrophages in HFpEF and control hearts.
12. **Supplement Table 1**. List of antibodies used for flow cytometry.
13. **Supplement able 2**. The parameters of mice at baseline and 5 weeks after high-fat diet and L-NAME.
14. **Supplement Table 3.** The functional roles of 9 causal genes.
15. **References.**

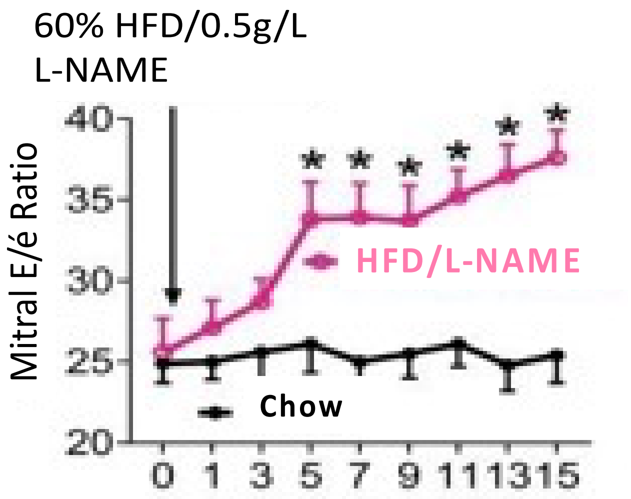


**Supplement Figure 1. Successful induction of HFpEF by HFD and L-NAME**. C57BL/6 mice at 12-14 weeks of age were fed with 60% HFD and 0.5 g/L L-NAME or normal chow as a control. The left ventricle (LV) was dynamically examined with a Visualsonics Vevo 3100 echocardiography. Mitral E/e' ratio (the index of diastolic function) was significantly elevated from 5 to 15 weeks after HFD and L-NAME feeding. Data are represented as mean ± SEM. *p<0.05, and Student's t test.

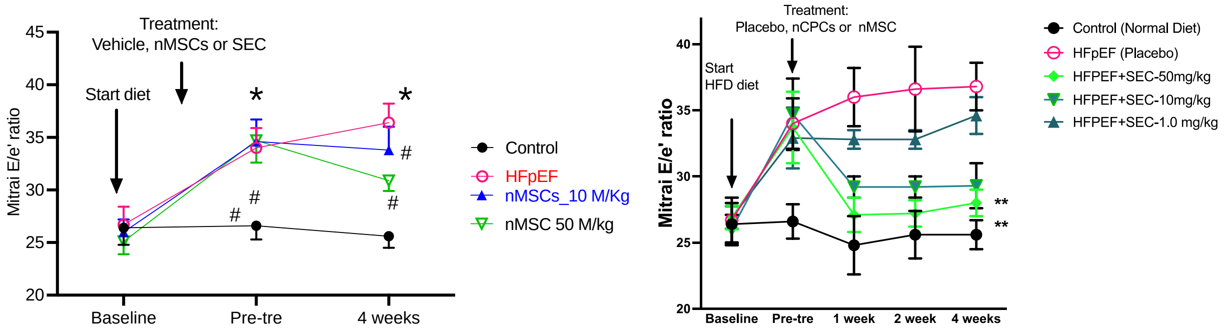


**Supplement Figure 2.** Optimal dosing of nMSCs or their secretome (SEC) for treating HFpEF. Mitral E/e' ratio (the index of diastolic function) was calculated after: a) administering either 10 M cells/kg or 50M cells/kg of nMSC (left) and b) three different doses of secretome (1.0 mg/kg or 10mg/kg or 50mg/kg). Data are represented as mean ± SEM. *, $p<0.05$, **, $p<0.01$, #, non-significant (ns), and analyzed using Prism Software multiple comparisons.

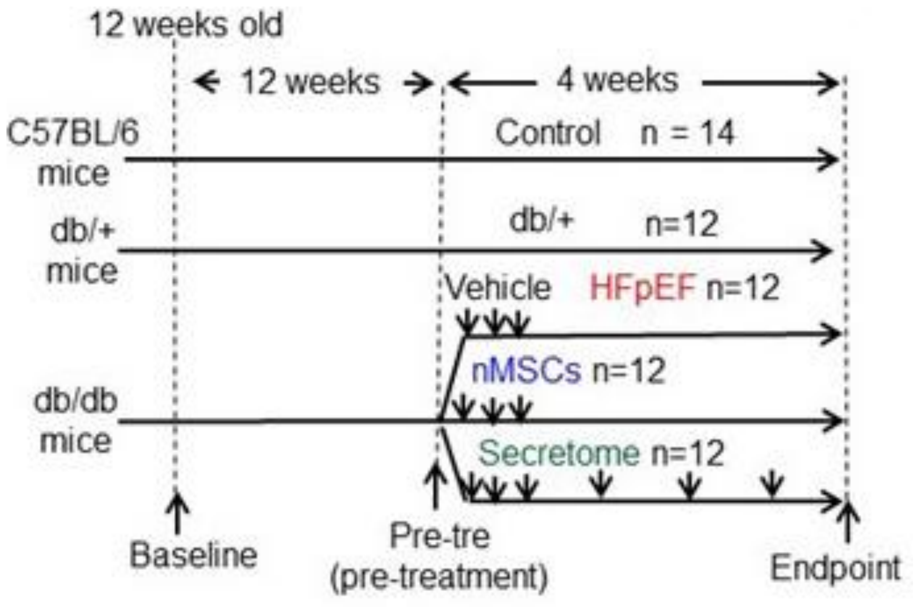


**Supplement Figure 3.** Schematic Study Design in db/db mice.

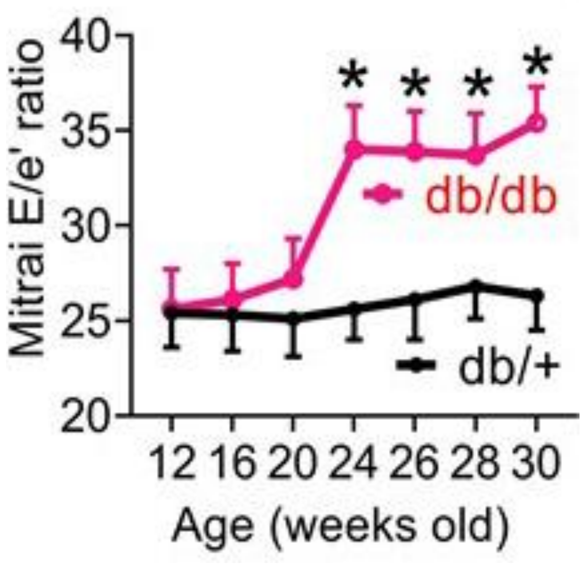


**Supplement Figure 4.** Db/db mice at 26 to 30 weeks of developed cardiac diastolic dysfunction. The left ventricle (LV) was dynamically examined with a Visualsonics Vevo 3100 echocardiography every week to assess the induction of HFpEF. The mitral E/e' ratio (the index of diastolic function) was significantly elevated from 20 to 24 weeks after inducing HFpEF in db/db mice. Data are represented as mean ± SEM. *$p < 0.05$ versus db/+ group, and analyzed by Student's t test.

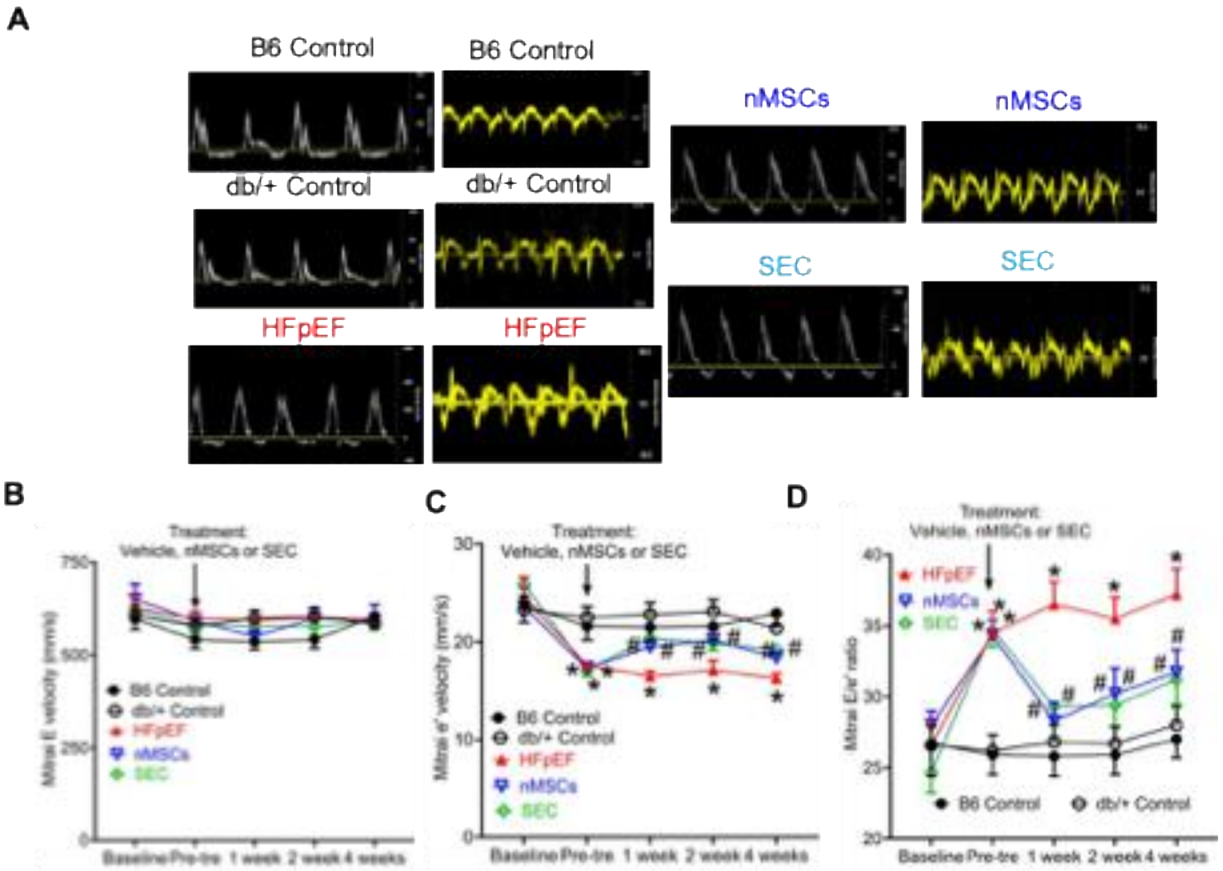


**Supplement Figure 5. Therapeutic efficacy in diastolic function of nMSCs or SEC in HFpEF db/db and db/+ mice.** Improved cardiac diastolic function by nMSCs and secretome (SEC) in the mice with HFpEF induced by HFD and L-NAME. A) Representative pulse Doppler and tissue Doppler waves showing mitral E and e’ waves. B) Mitral E velocity. C) Mitral e’ velocity. D) Mitral E/e’ ratio. *P<0.05 versus control, and #P<0.05 versus HFpEF groups (n = 12-14 mice/group).

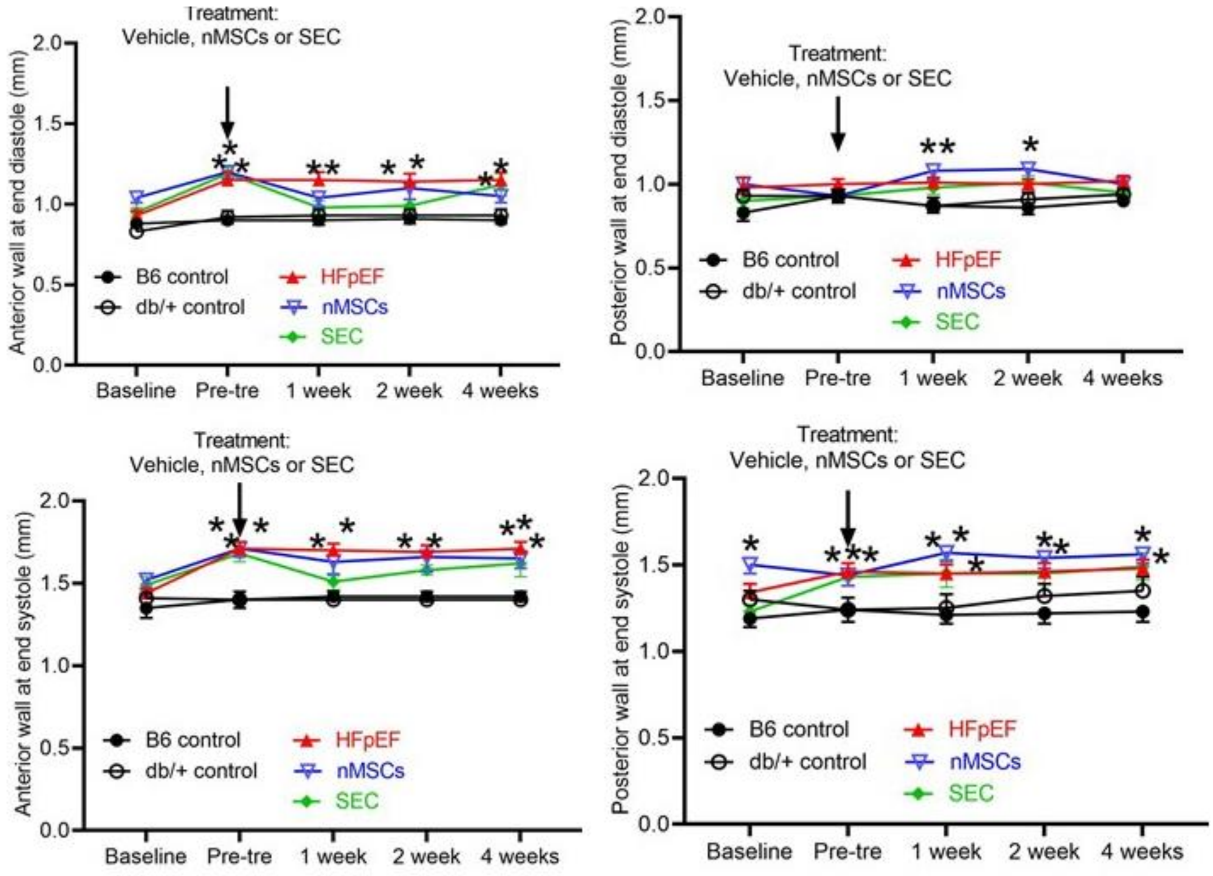


**Supplement Figure 6. Both nMSCs and secretome (SEC) did not change the wall thickness of the left ventricle in db/db mice.** The parameters of the left ventricular wall thickness were derived from short axis B-mode-guided M-mode images at baseline, pretreatment, and 4 weeks post treatment from C57BL/6 mice treated with PBS (B6 control), db/+ mice with PBS (db/+ control), and db/db mice with PBS (HFpEF), with nMSCs (nMSCs), or with secretome (SEC). Data are represented as mean ± SEM. *$p < 0.05$ versus B6 control and db/+ control, and analyzed by 1-way ANOVA followed by Tukey test.

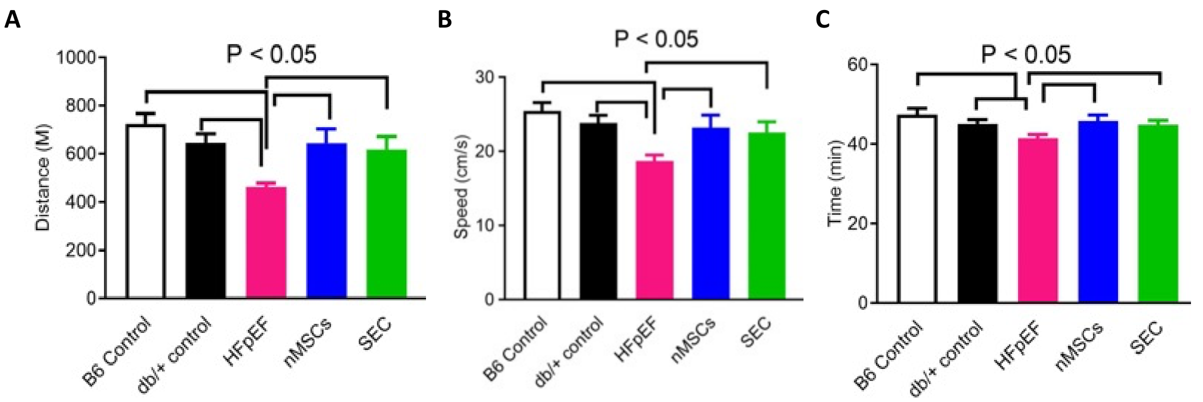


**Supplement Figure 7. Effect of nMSC and SEC on exercise tolerance of db/db induced HFpEF mouse model.** Significantly improved physical function was observed after treatment with nMSCs and SEC as compared to placebo treated HFpEF mice as determined by distance covered (A), speed (B), and time taken (C) for the distance. Data are represented as mean ± SEM (n = 12 – 14 mice/group)and were analyzed by one-way ANOVA followed by Bonferroni's test.

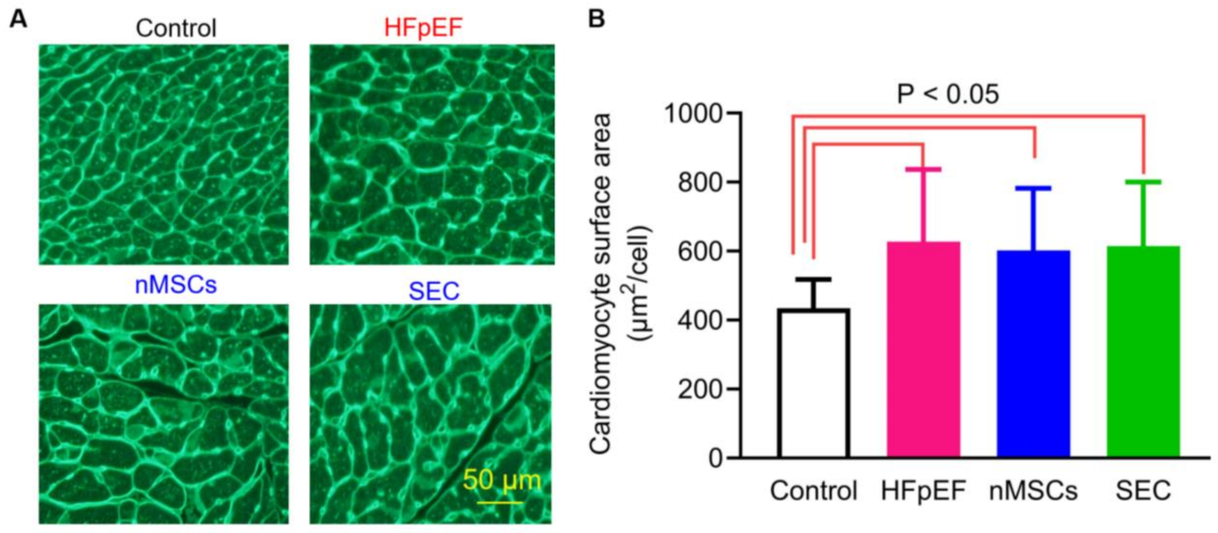

**Supplement Figure 8. Both nMSCs and SEC did not significantly change increased cardiomyocyte size in HFpEF mice.** A: Representative micrographs of heart sections of control, HFpEF, nMSCs, and secretome (SEC) mice stained with wheat germ agglutinin (WGA). Scale bar: 50 µm. B: Quantification of cardiomyocyte surface area in 4 groups of mice (n = 10 sections in 3 mouse hearts/group). Cardiomyocyte size was measured by staining of mouse hearts with WGA. Data are presented as means ± SEM. Kruskal-Wallis test followed by Dunn's test was used to analyze multiple group comparisons.

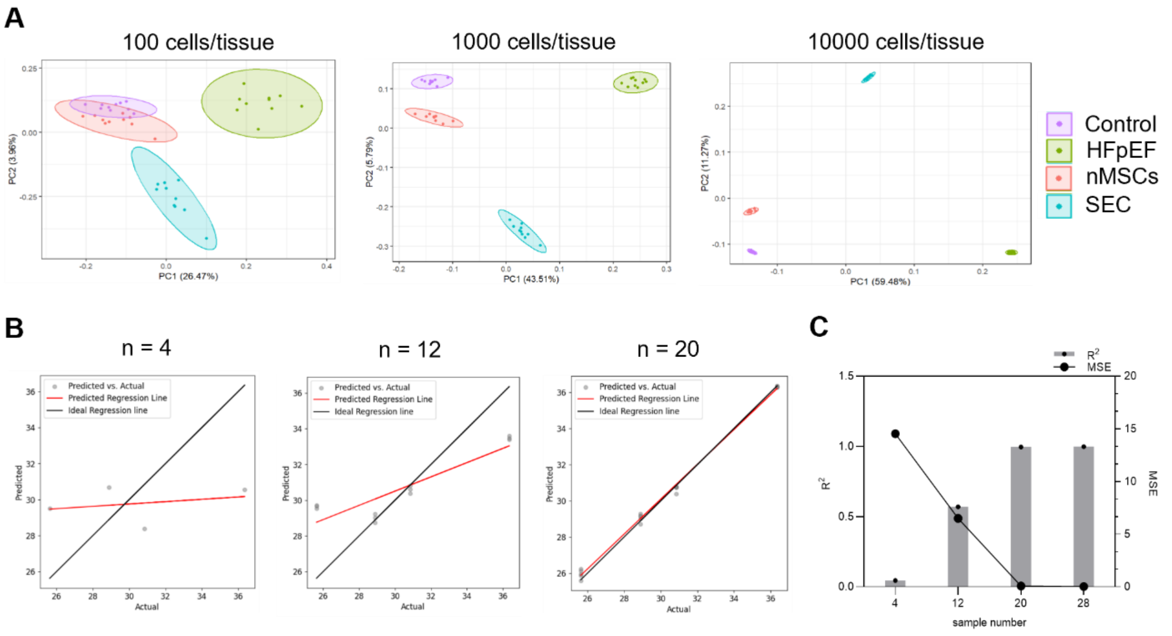


**Supplementary Figure 9. VIPcell framework enabled supervised and causal machine learning on limited number of snRNAseq data for measured tissue-level diastolic function.** A: Principal Component Analysis (PCA) on augmented bulkRNA-seq data. The analysis demonstrates clear separation between groups, with the number of bootstrapped cells per snRNA-seq datum influencing the clustering patterns. (n = 40, purple; Control, Green: HFpEF, Red; nMSCs, Blue; SEC) B: PLS regression prediction accuracy **c**, Scatter plots compare actual(x-axis) versus predicted(y-axis) values of E/E' across PLS regression models trained on varying numbers of augmented sample data. C: Model performance metrics as a function of augmented sample size. $R^2$; left y-axis, MSE; right y-axis.

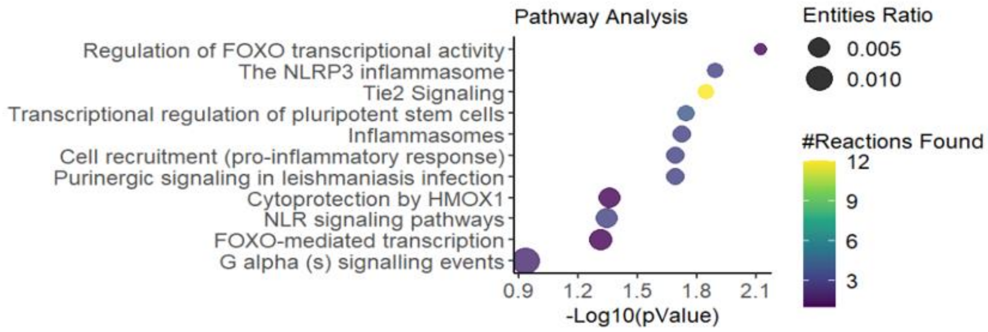


**Supplement Figure 10. Pathway enrichment analysis of nine direct causal genes for E/e' diastolic function identified from nMSCs- and SEC-treated HFpEF groups.** The bubble plot represents enriched pathways based on the Reactome database. The x-axis shows the -Log10(pValue), indicating statistical significance, while the y-axis lists the identified pathways. The size of the bubbles corresponds to the entities ratio, where larger bubbles indicate a higher proportion of genes associated with each pathway relative to the total detected genes. The color gradient represents the number of reactions found within each pathway, ranging from purple (fewer reactions) to yellow (more reactions).

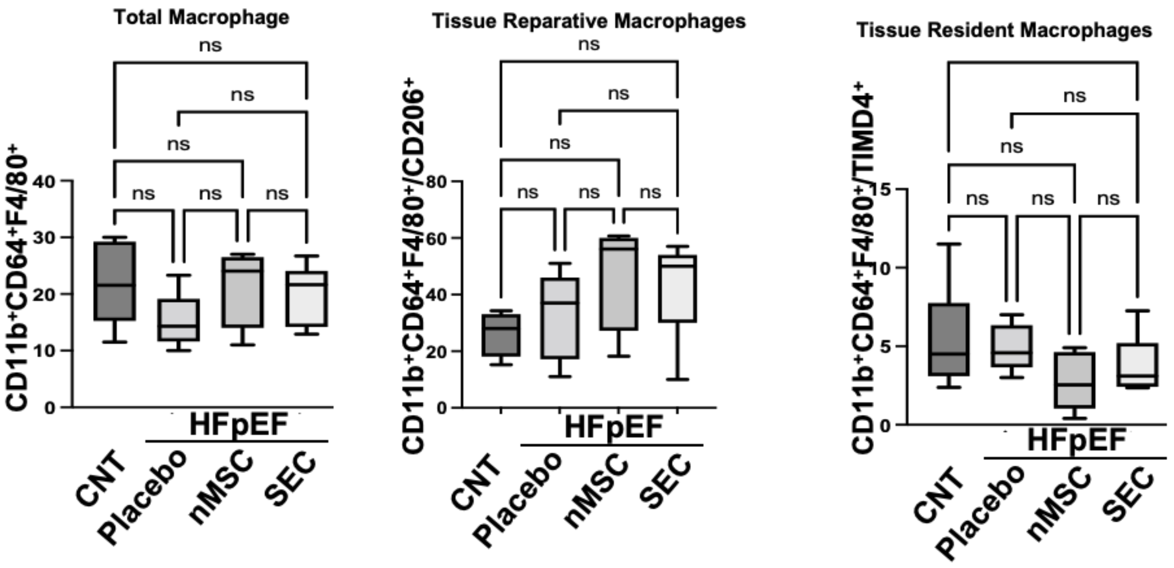


**Supplement Figure 11. Effect of nMSCs- or SEC-treatment on various types of macrophages in HFpEF and control hearts.** Flow cytometric analysis of Total Macrophage $CD11b^+CD64^+F4/80$, Tissue Reparative Macrophages $CD11b^+CD64^+F4/80^{+/}CD206^+$, and Tissue Resident Macrophages $CD11b^+CD64^+F4/80^{+/}TIMD4^+$(n = 5-6 hearts/group). Data was analyzed by Ordinary one-way ANOVA using Prism software (V 10). Holm-Sidak’s test was used for multiple comparisons. * p<0.05, **p<0.01, ns = not significant

**Supplementary Table 1.** List of antibodies used for flow cytometry studies

| No. | Primary antibodies | Supplier | Product No. |
|---|---|---|---|
| 1 | CD3 | BD Pharmingen (APC) | 557030 |
| 2 | CD4 | Biolegend (PE/Cy7) | 201516 |
| 3 | CD8 | BD Horozon (V450) | 561614 |
| 4 | CD11b | Invitrogen (FITC) | 12-0112-82 |
| 5 | CD19 | Invitrogen (APC) | 47-019842 |
| 6 | CD64 | Invitrogen (perCP-eFLuor719) | 46-0641-82 |
| 7 | F4/80 | Invitrogen (APC) | 17-4801-82 |
| 8 | FoxP3 | Invitrogen (PE) | 12-5773-82 |
| 9 | TIMD4 | Invitrogen (Alexa) | 53-5866-82 |

**Supplementary Table 2.** Characteristics of C57BL/6 mice fed high-fat diet (HFD) and L-$N^G$-nitroarginine methyl ester (L-NAME) water before (baseline) and after 5 weeks of diet

| | Baseline | | 5 weeks | |
|---|---|---|---|---|
| | Chow | HFD+L-NAME | Chow | HFD+L-NAME |
| Body weight (g) | 22.0±0.1 | 21.7±0.1 | 25.6±0.1 | 29.6±0.2* |
| Heart rate (beats/min) | 496±8 | 482±8 | 507±14 | 495±13 |
| Anterior wall at end diastole (mm) | 0.76±0.02 | 0.78±0.03 | 0.82±0.02 | 0.91±0.02* |
| Anterior wall at end systole (mm) | 1.22±0.04 | 1.24±0.05 | 1.29±0.04 | 1.42±0.03* |
| Posterior wall at end diastole (mm) | 0.77±0.03 | 0.75±0.02 | 0.82±0.03 | 0.93±0.04* |
| Posterior wall at end systole (mm) | 1.07±0.03 | 1.07±0.03 | 1.12±0.04 | 1.26±0.06* |
| LV internal diameter at end diastole (mm) | 3.81±0.06 | 3.78±0.06 | 3.98±0.07 | 3.99±0.07 |
| LV internal diameter at end systole (mm) | 2.59±0.08 | 2.50±0.08 | 2.77±0.09 | 2.78±0.09 |
| Fractional shortening (%) | 32±1 | 34±1 | 31±1 | 31±1 |
| LV end-diastolic volume (μL) | 63±2 | 62±2 | 70±3 | 70±3 |
| LV end-systolic volume (μL) | 25±2 | 23±2 | 27±2 | 28±2 |
| Ejection fraction (%) | 61±2 | 63±2 | 62±1 | 60±1 |
| LV mass (mg) | 82±3 | 81±3 | 97±4 | 19±1 |
| Cardiac output (mL/min) | 19±1 | 19±1 | 20±1 | 31±1 |
| Mitral E/e' ratio | 26±1 | 27±2 | 25±2 | 35±2* |

**Supplementary Table 3. The functional roles of 9 causal genes.**

| No. | Gene | Functional Roles | Ref. |
|---|---|---|---|
| 1 | Foxp1 | Enforcing Foxp3-mediated regulation of gene expression and enabling efficient IL-2 signaling in $T_{reg}$ cells | [4] |
| 2 | Arl15 | Encoding a small GTP-binding protein, involved in Tgf-β signaling, positively correlated with a CD4+ T cell | [5, 6] |
| 3 | Tmtc1 | Involved in cell proliferation and inflammation, and development of ER | [7] |
| 4 | Camk1d | Key modulator of immune resistance, co-expressed with PD-L1 | [8] |
| 5 | Angpt1 | Involved in angiopoietin/TIE2 pathway, increasing the infiltration of T cells | [9, 10] |
| 6 | Txnip | Critical metabolic regulator of $T_{reg}$ identity and function | [11, 12] |
| 7 | Pde3a | Regulating the levels of cyclic adenosine monophosphate (cAMP) and cyclic guanosine monophosphate (cGMP) in cells, Enriching Foxp3+ T cell | [13, 14] |
| 8 | Stox2 | Preventing glioblastoma stem cells from being recognized by the immune system | [15] |
| 9 | Etl4 | Expressed in the notochord of early embryos and in multiple epithelia during later development | [16] |
